# Offline Memory Reactivation Promotes the Consolidation Of Spatially Unbiased Long-Term Cognitive Maps

**DOI:** 10.1101/2020.08.20.259879

**Authors:** Andres D. Grosmark, Fraser T. Sparks, Matt J. Davis, Attila Losonczy

## Abstract

Spatial memories which can last a lifetime are thought to be encoded during ‘online’ periods of exploration and subsequently consolidated into stable cognitive maps through their ‘offline’ reactivation^1–5^. However, the mechanisms and computational principles by which offline reactivation stabilize long-lasting spatial representations remain poorly understood. Here we employed simultaneous fast calcium imaging and electrophysiology to track hippocampal place cells over weeks of online spatial reward learning behavior and offline resting. We describe that recruitment to persistent network-level offline reactivation of spatial experiences predicts cells’ future multi-day representational stability. Moreover, while representations of reward-adjacent locations are generally more stable across days, reactivation-related stabilization is, conversely, most prominent for reward-distal locations. Thus, while occurring on millisecond time-scales, offline reactivation counter-balances the observed multi-day representational reward-adjacency bias, promoting the stabilization of cognitive maps which comprehensively reflect entire underlying spatial contexts. These findings suggest that post-learning offline-related memory consolidation plays complimentary and computationally distinct role in learning as compared to online encoding.

---

Cognitive maps can support flexible problem solving, such as finding novel paths to and from rewards, which otherwise pose challenges to purely operant-based learning strategies ^6^. In order to be useful, such maps need to comprehensively represent the environment, rather than just the paths the animal takes most frequently or those associated with rewards. The encoding and consolidation of the hippocampal-dependent cognitive maps involves neural mechanisms operating across a wide range of time-scales, from milliseconds to days. During exploratory (‘online’) periods a 5-10 Hz (theta) oscillation observed in the local field potential (LFP) of rodents temporally organizes the spatial receptive fields of hippocampal place cells ^7^, which themselves show a bias towards over-representation of rewarded locations ^8^. Conversely, during quiet resting (‘offline’ periods) hippocampal activity is organized into short (50 to 100 ms) sharp-wave/ripple (SWR) events which are associated with transient 125-225Hz LFP oscillations and the time-compressed replay of previously encoded hippocampal representations ^9–11^. SWR-related reactivation is in turn thought to support the offline consolidation of hippocampal-dependent memory traces ^2,3,12^. However, much remains unknown regarding the long-term persistence of memory replay, its effects on subsequent online spatial representations and its relationship to the motivational gradients which drive spatial behaviors.

Calcium imaging techniques have revealed that, similar to the partial degradation of online spatial memory performance across days, longitudinal hippocampal spatial activity is characterized by partial and decaying stability ^13,14^. However, the absence of well-defined calcium-based biomarkers of offline reactivation have so far limited attempts to uncover the contribution of offline consolidation to the stabilization of memories over the long behavioral timescales over which they are expressed. Here, we track the formation, consolidation and re-expression of hippocampal spatial memory traces across time-scales. We report that hippocampal spatial memory traces show persistent offline SWR-related reactivation and that these post-learning reactivations predict the stability of future online spatial coding. Moreover, we find that while reward-adjacent spatial representations show overall enriched multi-day stability, this reward-bias is counterbalanced by SWR-related offline stabilization which is most prominent for representations farther from the reward. Therefore, we propose that post-learning offline memory consolidation promotes the formation of spatially comprehensive, as opposed to narrowly reward-biased, cognitive maps of space.

We performed 60 Hz two-photon calcium imaging of genetically identified CA1 pyramidal cells with simultaneous contralateral CA1 LFP recordings (Fig. 1A) in the dorsal hippocampus of head-fixed CaMKII-Cre mice (n = 10). The same field of view and (FOV) and identified cells were stably imaged over 12 days (Fig. 1B-C; fig. S1) allowing us to track the long-term evolution of place coding. Each day the mice were recorded during 3 consecutive 15 to 20 minute sessions. PRE and POST imaging sessions were performed on a cue-less and un-rewarded burlap belt (belt A) on which the animal typically rested quietly, while during the intervening RUN session the mice ran for a spatially stable operant reward on one of two distinct 2-meter cue-rich treadmill belts (belts B and C, Fig. 1C), allowing us to test the specificity of the observed spatial memory traces and their relationship to a spatially located goal. As expected, during RUN sessions the animals’ behavior was strongly biased by the reward such that they spent the most time near, and ran fastest away from, the rewarded location (Fig. 1D). As mice ran through the environment the calcium-estimated spiking activity of CA1 pyramidal cells (fig. S2) showed robust spatial coding, while the LFP oscillated at theta frequency (Fig. 1E). The calcium activity of individual place cells was strongly modulated by the ongoing theta oscillation (Fig. 1F; fig. S3), confirming that two-photon calcium imaging has sufficient temporal resolution for resolving fine-timescale network dynamics during the online state. Individual place fields were specific to a given RUN belt, and showed a spatially differential decaying level of stability across days (Fig. 1G-I). Notably, place cells with coding peaks near the reward tended to show relative cross-day coding stability as compared to those coding for locations further from the reward (Fig. 1J; figs. S4-5). Bayesian decoding of spatial location within and across days further confirmed the context-specificity and partial multi-day decay of the population-level hippocampal spatial code (Fig. 1K-M; fig. S6). Together these findings confirm that hippocampal spatial coding is characterized by both stable and unstable multi-day coding elements.

**Figure 1:**
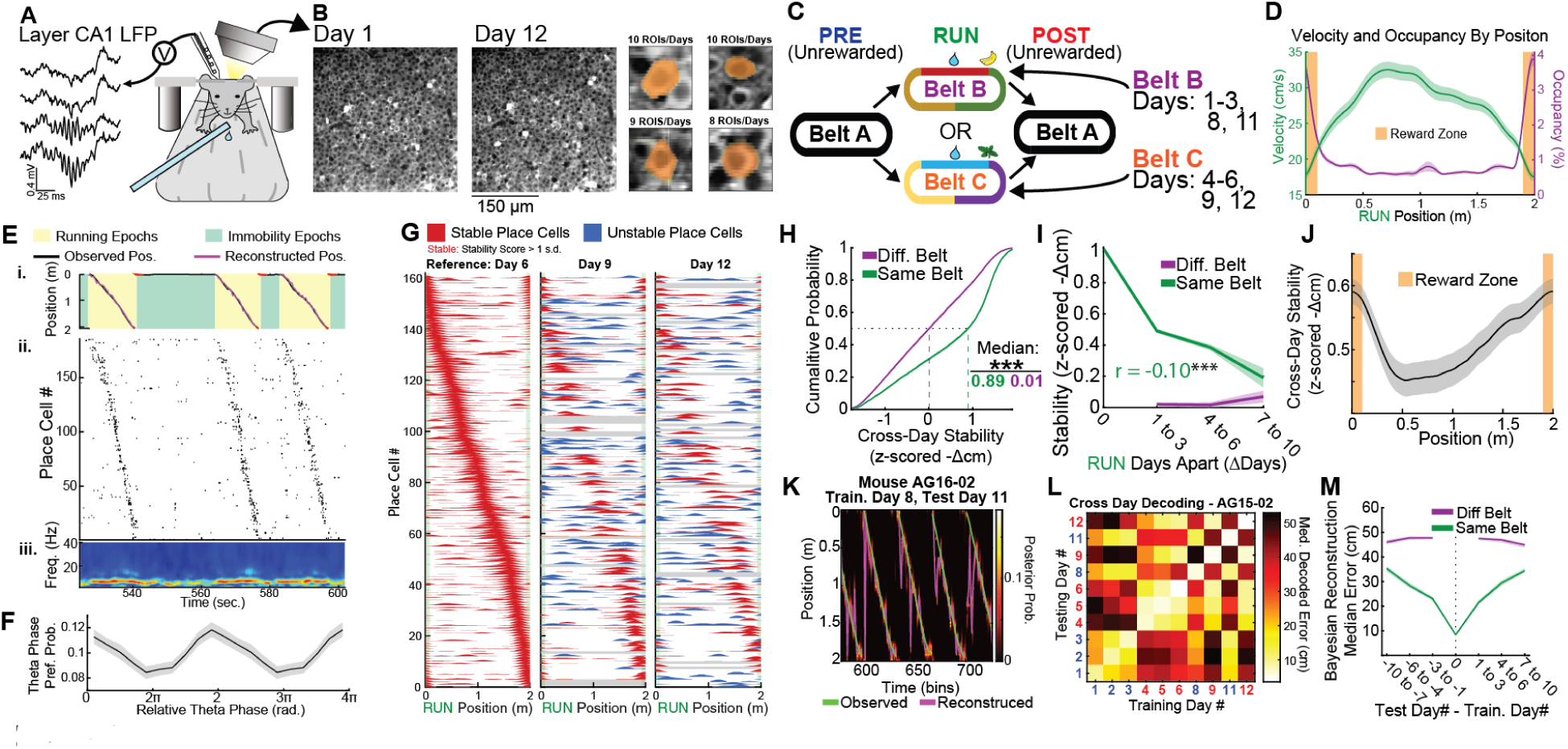
Long-term calcium-imaging and LFP recordings reveal both stable and unstable long-term hippocampal spatial coding. **(A)** Simultaneous CA1 pyramidal layer two-photon calcium imaging and contralateral CA1 LFP recordings were carried out on head-fixed mice. **(B)** Populations of genetically identified CA1 pyramidal cells were stably imaged throughout the duration of the experiment (right panel shows 4 example cross-day registered ROIs). **(D)** Each day the mice were imaged/recorded during consecutive PRE, RUN, and POST sessions. During each RUN the animals performed a hidden spatial reward task on one of two cue-rich belts (belts B and C). (**D**) Mice ran slower (left-axis) and spent the most amount of time (right-axis) near the rewarded locations (orange, graphs show the mean ±SEM in a sliding 20 cm window). (**E**) An example of three RUN laps showing (i) the position of the animal (black line) could be decoded (purple line) using (ii) the deconvolved calcium activity of the 180 place cells simultaneously imaged during this session. (iii) The wavelet spectrogram of the CA1 LFP showed increases in theta-band power during locomotion. (**F**) Population histogram of relative theta-phase preference across all place cells shows robust LFP theta-phase related modulation of the imaged place cell activity (p ~ 0, Rayleigh test). (**G**) Example from place cells imaged across 3 days on Belt C arranged by their peak firing location on day 6 (left panel). Cells with stable place coding (stability score > 1) relative to day 6 are shown in red, unstable coding place cells are shown in blue (this threshold is used only for visualizing this example). Note the simultaneous presence of both stable and unstable multi-day spatial coding elements. (**H**) Cumulative distribution of cross-day stability scores across different (purple) or same (green) RUN belts (same-belt median score = 0.89, n = 9,721 cell-pairs, different-belt median = 0.01, n = 11,134 cell-pairs, p ~ 0, Ranked-Sum test). (**I**) Spatial coding stability decays over time (0 days apart compares first versus second half of RUN session, same-belt Pearson’s r = −0.10, p~0). (**J**) Spatial coding near the reward location (orange) is more stable over days (graphs show mean ±SEM of cross-day stability score in a rolling 20 cm bin across the belt, Pearson’s correlation between cross-day stability and place field peak distance to reward: r = −0.042, p < 3.84×10^−5^). (**K**) Example of cross-day Bayesian location decoding in which place fields templates from day 8 RUN are used to decode the animal’s location on day 11 RUN. (**L**) Error matrix showing the median location decoding error across days, belt B and C sessions are labeled blue and red respectively. (**M**) Group data of median across and within-day decoding error for different and same belt comparisons (same-belt cross-day median: 28.1 cm, n=11,320 laps, different-belt median: 47.0 cm, n=14,436 laps, Ranked-sum p ~ 0). (p* < 0.05, p** < 0.005, and p***< 0.0005 for all figures).

Next we examined SWR events occurring during PRE and POST immobility to assess place cell recruitment and learning-related changes in offline reactivation content. LFP-detected SWR events were associated with significant increases in place cell firing rates as indicated by their SWR-firing rate gain (within-SWR firing rate normalized by overall immobility firing rates, Fig. 2A,B, ^15^). Conversely, calcium-detected population synchrony events (PSE’s) in which at least 5 place cells became co-active within a short time-window during immobility were associated with increases in LFP-detected SWR-frequency LFP power (Fig. 2C). These results confirm the accessibility of short-time scale offline synchronous activity to our joint calcium-imaging/LFP approach. Consistent with the learning-dependence of SWR-dynamics ^16^, SWR incidence, duration, and place cell recruitment increased POST as compared to PRE the RUN sessions (Fig. 2D, figs. S7-8). Moreover, PRE to POST increases in pair-wise synchrony (firing rate correlations) were not uniform, but were largest for pairs of place cells with nearby place field peaks during the intervening RUN (Fig. 2E; fig. S9), consistent with the reactivation of RUN-related patterns. We further examined this reactivation by estimating ensemble ‘templates’ of co-active neurons during the RUN (‘RUN-ensembles’) using independent component analysis (ICA) and calculating their reactivation during offline states ^17,18^. RUN-ensembles were found to be more strongly reactivated during the POST as compared to PRE sessions, an effect which was largest around the time of SWRs (Fig. 2F and G, respectively; fig. S10, A-F). Having confirmed the presence of within-day ensemble reactivation, we then analyzed the within-day PRE to POST reactivation changes on a given day (test day) for RUN-ensemble templates calculated on a different day (train day). While smaller than within-day changes, significantly larger PRE to POST increases in reactivation were observed for pairs of days in which the animal ran on the same belt than for pairs on different belts (Fig. 2H), confirming the context specificity of the reactivated ensembles. Moreover, we observed that RUN-ensembles computed on a given day better explained the peri-SWR covariance of the subsequent day’s PRE epochs (~24 hours after the RUN) than of the same day’s PRE epochs (~20 minutes before the RUN), but thereafter fell to chance levels, demonstrating the multi-day time-course of the offline reactivation of specific hippocampal spatial ensembles (Fig. 2I; fig. S10, G-Q). Together, these results demonstrate the long-term dynamics of the offline reactivation of context-specific RUN-related neural ensembles.

**Figure 2:**
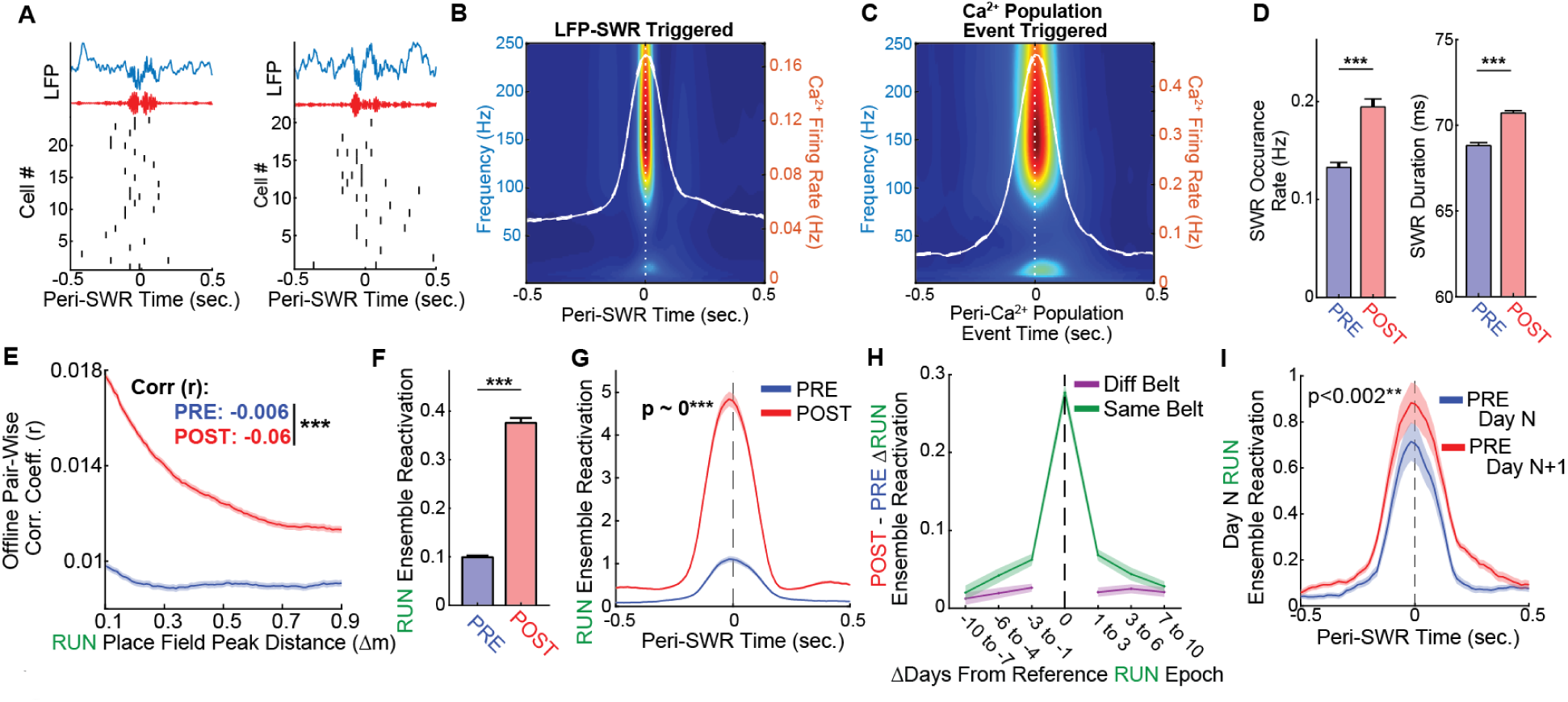
Context Specific and Long-Lasting Offline Reactivation of RUN Related Neural Ensembles. (**A**) Two example SWR’s showing the peri-SWR LFP (blue), SWR-filtered LFP (red), and deconvolved calcium activity of the cells active within each SWR event (black spike rasters). (**B**) Average peri-SWR deconvolved firing rate (right-axis, foreground white-lines, mean ±SEM) overlaid on the average peri-SWR wavelet spectrogram (heat-map, left axis). (**C**) Average population synchrony events (PSEs) detected in the calcium-activity (plotted as in *B*). (**D**) SWR events were more frequent (left panel) and of longer duration (right panel) during POST as compared to PRE epoch immobility (panels show mean ±SEM). (**E**) Pairs of place cells that coded for nearby locations on the RUN belt (x-axis) showed elevated pair-wise rate correlations during the POST as compared to PRE epochs (y-axis, p ~ 0, Fisher Z-test, n=1,144,357 place cell pairs). (**F**) ICA ensembles computed during RUN were preferentially reactivated in the POST as compared to the PRE epoch (p ~ 0, Signed Rank test, n = 3,844 components). (**G**) Preferential POST epoch reactivation of RUN ensembles was most pronounced around the time of SWR events (p ~ 0, Signed Rank test). (**H**) Cross-day RUN ensemble PRE to POST changes in reactivation are greater for same-belt than different-belt cross-day pairs (same-belt median reactivation = 0.0187, n = 8,603 components, different-belt median reactivation = 0.009, n = 12,915 components, p ~ 0, Ranked Sum test). (**I**) RUN ensembles showed significantly greater reactivation during SWR-events during the PRE on the day following a RUN (~24 hours after a RUN, red lines, means ±SEM) than during the PRE immediately before the RUN (~20 minutes prior a RUN, blue lines, p < 0.002, Signed-Rank test, n = 1,750 components).

We next asked whether sequential spatial representations are reinstated in the fine time-scale calcium activity of place cells during offline PSEs, and if so, how their content changed after learning. The spatial content of individual PSE’s was estimated from the Bayesian decoding of calcium-estimated spikes occurring in two-frame (~33ms) bins (Fig. 3A, ^19^). We found that individual PSE’s significantly replay RUN spatial activity sequences measured in both the forward and reverse direction as compared to bin-shuffled events (Fig. 3B, ^20–22^). Significantly more sequence replay content was observed during RUN and POST PSE’s than during the PRE PSE’s (Fig. 3C-D; ^16^), and these differences were robust across a variety of computational controls (fig. S11), confirming that the sequence replay of both local and distal environments is accessible through two-photon calcium imaging techniques. Moreover, not only did the presence of spatial sequences increase during and after the RUN, but the spatial content of significant replay events changed as well, with POST epoch replay events coding for locations farther from the reward than PRE events (Fig. 3E-F). Likewise, the decoded spatial content across all PSE’s was found to change systematically across epochs (Fig. 3G-H). Particularly, we found that while spatial content was biased towards the rewarded location across all epochs, POST epoch PSEs decoded for locations further from the reward than did PRE or RUN epoch PSE’s. This pattern of encoded locations was confirmed for SWR-related activity (Fig 3I). Therefore, while overall decoded activity remains biased towards the rewarded location, this bias is diminished by the putatively learning-related changes in activity observed from PRE to POST. Notably a comparison of decoded content during SWR and non-SWR immobility periods revealed that SWR content was significantly less biased towards the rewarded location than non-SWR content (Fig. 3J), suggesting that SWRs are privileged windows for the relatively homogenous reactivation of entire spatial environments, rather than only their most salient portions.

**Figure 3:**
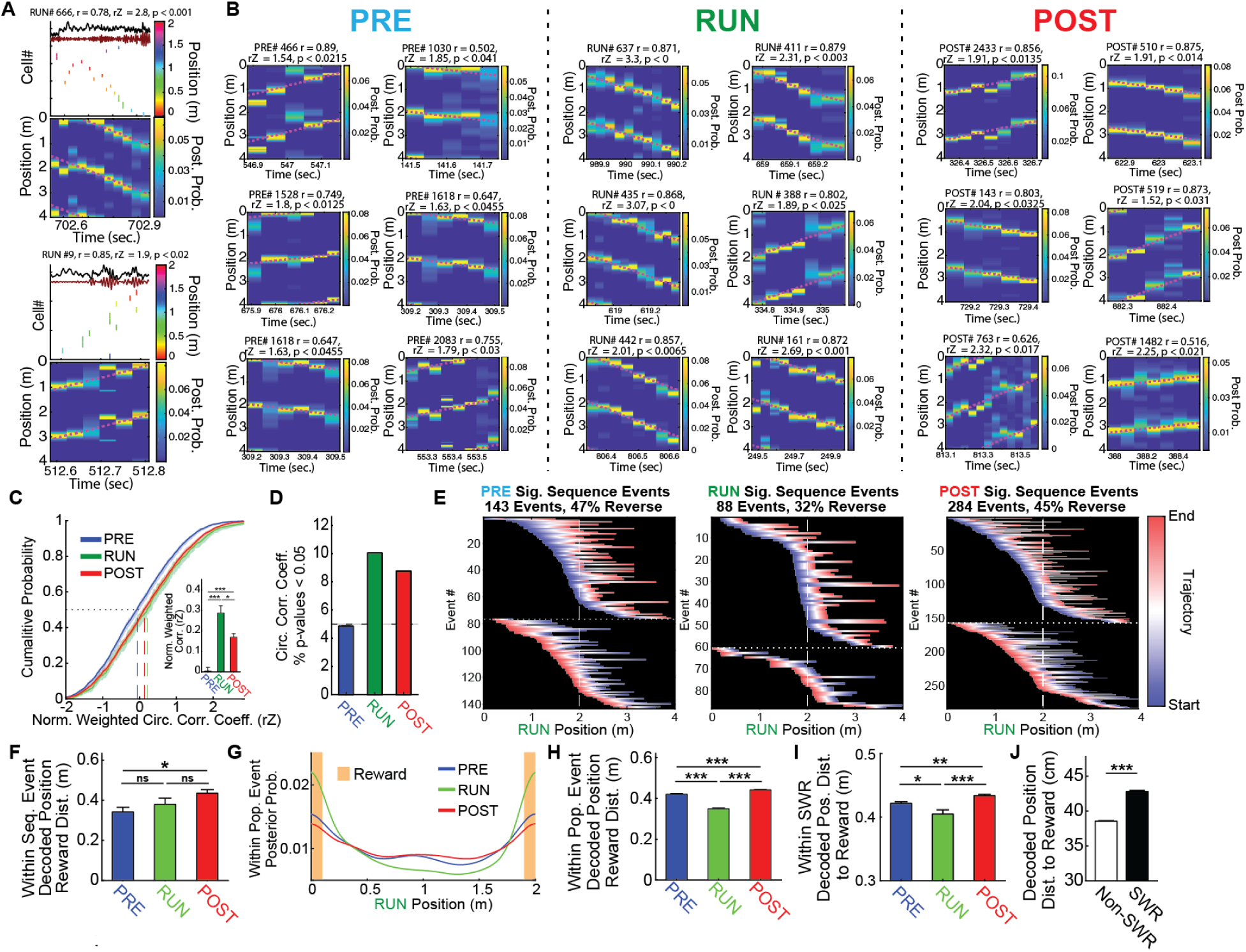
Post-Learning Place Sequence Replay is Biased Away from Rewarded Locations. (**A**) Two example RUN epoch PSE’s (occurring during immobility) with significant forward (top) or reverse (bottom) replay are shown, including the raw and SWR-filtered LFP traces (black and red lines), the raster of the participating place cells sorted by place field position, and the Bayesian decoded posterior probability of position (tiled twice to account for reward-spanning trajectories). (**B**) Six example significant sequence PSE’s are shown for the PRE, RUN and POST epochs (magenta lines show the best fit trajectories). (**C**) Cumulative distribution of shuffle-normalized sequence scores (rZ) across behavioral epochs. Significantly more sequence content was observed during the RUN and POST epochs as compared to the PRE epoch (inset, mean ±SEM, Kruskal-Wallis test p < 7.7×10^−14^, Post-Hoc Tukey tests). (**D**) Percentage of PSE’s with significant (p < 0.05) sequence content relative to temporal-bin shuffle during each behavioral epoch (143/2,978 PRE, 88/993 RUN, 284/3,438 POST PSEs). (**E**) Heat maps of the linearized best-fit line occupancy of significant trajectory events by behavioral epoch (blue and red show event start and end, respectively, horizontal line separates forward (top) from reverse events (bottom), position is tiled vertically twice to account for reward-spanning sequences, the single rewarded location corresponds to positions 0m, 2m and 4m on the maps). (**F**) Significant sequence PSEs during the POST code (center of mass) for locations further than PRE significant sequence PSEs (Kruskal-Wallis test p < 0.008, followed by post-hoc Tukey test). (**G**) Bayesian reconstructions of RUN belt location within PSEs shows distinct reward-related biasing during different behavioral epochs (mean ±SEM). (**H**) POST epoch PSEs tend to encode for locations further from the reward than do PRE or RUN PSEs (reconstruction center of mass across PSEs, mean ±SEM, Kruskal-Wallis test p < 9.7×10^−19^, Post-Hoc Tukey tests). (**I**) POST epoch SWR content is less biased towards rewarded locations than PRE or RUN SWR-content (mean ±SEM, Kruskal-Wallis p < 8.1×10^−5^). (**J**) During immobility within-SWR reconstructed content is less biased towards the rewarded locations than non-SWR content (mean ±SEM, pooled across all behavioral epochs, Ranked-Sum p ~ 0).

We next examined the role of post-learning recruitment of SWRs on long-term spatial memory consolidation. For each day *N*, cells showing novel place fields were divided by session into two equal groups by their POST – PRE SWR-specific change in activity (SWR-Gain, ‘per cell’ analysis, see Online Methods; Fig. 4A). Day *N* RUN place cells displaying High SWR-Gain tended to more stably retain their place field firing locations onto future days on the same belt (Fig. 4B-C). This spatial stabilization effect was specific to SWR periods (Fig. 4D) and was not accounted for by spatial coding or rate differences between the Low and High SWR-Gain groups during the day *N* RUN epoch (fig. S12). Notably, an analysis of the SWR-associated stabilization revealed that this effect was not evenly distributed across the RUN belts, but instead was largest for locations further from the reward (Fig. 4E-F). In order to further confirm this observed spatial-bias of the SWR stabilization effect, the analysis above (Fig. 4B) was repeated for cells with day *N* place fields either nearer to (Goal zone cells), or further from (Path zone cells) the reward, excluding cells coding for the reward zone itself revealing that SWR-related stabilization was specific to Path zone cells (Fig. 4G). The interaction between place field distance to reward and SWR-related spatial stabilization was consistent across animals and 6 computational methods (fig. S13-14), and was further confirmed through an independent generalized linear modelling approach which also controlled for other day *N* place coding properties (fig. S15). A separate analysis in which all day *N* and day *N+i* same place cell pairs were considered (‘per-same-cell-pair’ analysis, see Online Methods) revealed that the path zone specific SWR-related stabilization effect is long lasting (> 7 days, Fig. 4H). Likewise, cross-day Bayesian reconstruction accuracy was improved in the High as compared to Low SWR-Gain cell place cell populations preferentially for areas of the belt further from the reward (Fig 4J; fig. S16). Finally, in an approach parallel to the SWR-Gain analysis above (Fig. 4B) day *N* place cells were divided by session into equal groups depending on their contribution to the PRE to POST changes in RUN-ensemble reactivation as estimated by a leave-one-out analysis. Place cells’ day *N* contributions to increased POST RUN-ensemble reactivation were predictive of their place coding stability onto future days, an effect which was found to be specific to Path zone coding place cells and absent in the Goal zone (Fig. 4K). Together, these results support the hypothesis that learning related PRE to POST changes in SWR-associated reactivation of place cells preferentially stabilizes the low-occupancy, high-velocity parts of the environment far from the reward.

**Figure 4:**
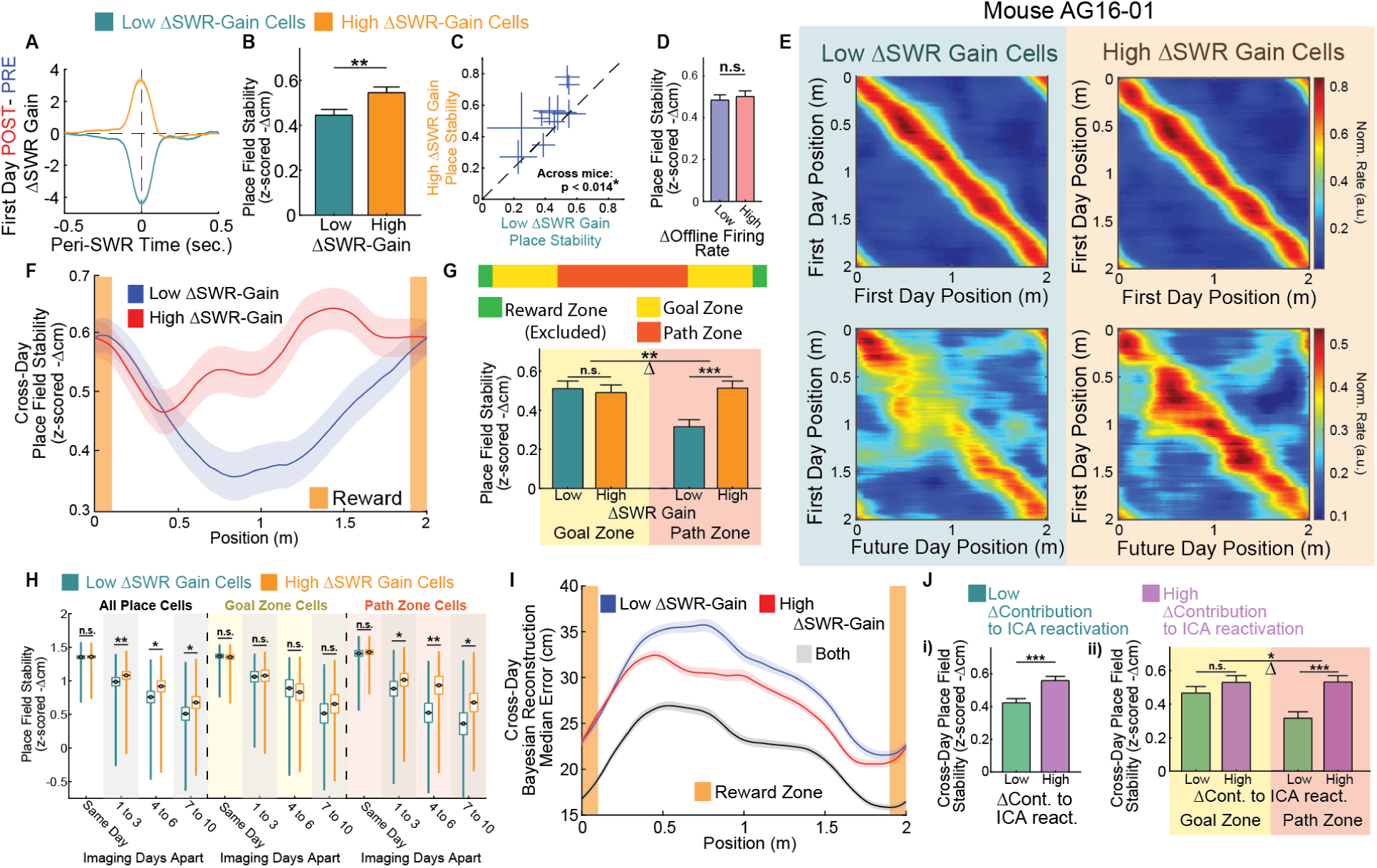
Learning related SWR-specific place cell recruitment predicts cross-day spatial stability. (**A**) For each day cells showing novel place fields were divided into two equal groups by their POST – PRE change in within-SWR gain (within-SWR firing rate normalized by total immobility firing rate, mean ±SEM, n = 1,212 cells per group). (**B**) PRE to POST increases in SWR-specific recruitment of novel place cells was predictive of their long-term spatial coding stability (mean ±SEM, Ranked-Sum test p < 0.004). (**C**) This effect was consistent across animals (crosses show by-animal mean ±SEM cross-day stability, Signed-Rank p < 0.014, n = 10 animals). (**D**) Conversely, general PRE to POST changes in immobility firing rates were not predictive of future spatial stability, confirming that the stability effect is SWR-specific (Ranked-Sum test, p < 0.44). (**E**) Place cells were sorted by their place field peak locations on the first day their place fields were expressed on a given RUN belt (first day, top panels) either for Low (left panels) or High (right panels) ΔSWR-Gain (place fields peaks were normalized to 1, plots show the mean sorted place field activity in a sliding 20 cm window). (**F**) High ΔSWR-Gain cells tended to show elevated cross-day coding stability specifically for low-occupancy, high-velocity locations farther from the reward (mean ±SEM). (**G**) In order to assess the interaction between distance to reward and putative SWR-related POST epoch stabilization, the RUN belt was divided into 3 areas (top panel): the reward area (20 cm) where the animal received the reward was excluded, the remained of the belt was divided equally into the goal zone (90 cm), near the reward, and the path zone (90 cm) farther from the reward. The analysis shown in (B) was repeated in these subgroups revealing that PRE to POST changes in SWR-Gain were predictive of future stability specifically for path zone coding place cells (Ranked-Sum test, p < 6.4×10-5, n=509 cells per group) and not goal zone cells (p < 0.79, n=513 cells per group) – and revealing a significant interaction between coding location and the effect of SWR-Gain changes in stability (Goal vs Path cell id permutation test for interaction, p < 0.0048). (**H**) In a complementary analysis all pairs of days for a which a given cell was a place cell on the same RUN belt were considered (within-cell pair-wise analysis). Place cells which showed High PRE to POST epoch changes in SWR-specific recruitment showed elevated spatial stability (y-axis) onto future days, an effect which was stable over time (x-axis) and specific to Path zone (right set of graphs, red shading) as opposed to goal zone (middle set of graphs, yellow shading) cells (Ranked-Sum tests, same day plots show stability between first and second half of laps within a day). (**I**) Bayesian reconstruction of position further confirmed that high ΔSWR-Gain cells display higher cross-day spatial coding stability preferentially at locations father from the reward (Goal vs Path cell id permutation test for interaction, p ~ 0). (**J**) A similar analysis as shown in B was performed based on each cell’s normalized PRE to POST change in contribution to ensemble (ICA) reactivation. Cell’s PRE to POST increases in contribution to ensemble reactivation were predictive of future place coding stability (panel i, Ranked-Sum test p < 1.5×10^−4^, n = 1,212 cells per group). Furthermore, this effect was specific to path zone as opposed to Goal zone cells (goal zone p < 0.19, Ranked-Sum test, Path zone cells p < 5.7×10^−5^ - Goal vs Path cell id permutation test for interaction, p < 0.03).

By combining fast two-photon calcium imaging and LFP recordings we tracked the formation and evolution of hippocampal memory traces across behavioral states. We demonstrate the presence of long-lasting, context specific replay and the association between post-learning SWR-specific recruitment and the subsequent spatial coding stability of individual place cells. Surprisingly, unlike the strongly reward biased statistics observed in the animal’s behavior and spatial map, the relationship between SWR-recruitment and spatial-coding stabilization was found to be largest in the low-occupancy, high-velocity areas of the environment further from the reward. These findings further support previous reports that the representational statistics of offline reactivation differ from those of online exploration ^23–25^. Specifically, consistent with the promotion of abstract rule or ‘gist’ learning by offline states observed in humans and rodents ^26,27^, hippocampal SWR-mediated memory consolidation may selectively promote the formation of integrated cognitive maps ^6,23,25^ as opposed to discrete reward-dependent behavioral programs ^28^.

## Materials and Methods

### Mice and viruses

All experiments were conducted in accordance with the NIH guidelines and with the approval of the Columbia University Institutional Animal Care and Use Committee. Imaging experiments were performed in 10 transgenic CaMKIIα-Cre mice on a C57/B16 background (R4Ag11 line, n = 4, Jackson Laboratory, Stock No: 027400, and R1Ag5 line, n = 6, Jackson Laboratory, Stock No: 027310, Dragatsis and Zeitlin, 2000). Cre-dependent recombinant adeno-associated virus (rAAV) expressing GCaMP6f (n = 6 mice, rAAV1-Syn-Flex-GCaMP6f-WPRE-SV40, Penn Vector Core) or GCamP7f (n = 4 mice, pGP-AAV-syn-FLEX-jGCaMP7f-WPRE, Add-Gene Catalog #104492-AAV1) were used for imaging of pyramidal neurons.

### Imaging window implant

Unilateral dorsal CA1 was stereotaxically injected with the rAAVs using a Nanoject syringe, as described previously ^30,31^. Injection coordinates were −2.2 mm AP, 1.75 mm ML relative to Bregma, and −1.2, −1.1, −1.0 mm DV relative to the cortical surface. 64 nL of diluted virus (1:12 in sterile cortex buffer) was injected at each DV location in 32 nL increments at a flow rate of 23 nL/sec. Three days later, mice were then surgically implanted with an imaging window (diameter: 3.0 mm; height: 1.5 mm) over the left dorsal CA1. Imaging cannulas were constructed by adhering a 3 mm glass coverslip (Edmond Optics, NJ) to a cylindrical stainless-steel cannula (3 mm diameter × 1.5 mm height) using optical adhesive (Norland, NJ). The surgical procedure was very similar to that previously described ^13,31,32^, the imaging window was implanted just above the alveus fibres, providing optical access to the pyramidal cells in the dorsal CA1. Briefly, following induction of anesthesia (Isoflurane: 3.5% induction, 1.5-2.0% maintenance; 1.0 L/min O_2_) and administration of analgesia (buprenorphine or meloxicam), the scalp was removed, and a 3.0 mm diameter craniotomy centred over the injection location was performed using a fin-tipped dental drill. The dura was removed, and the underlying cortex aspirated until fibers within the alveus were visible. The cannula with window was placed within the aspirated cavity and fixed to the skull with dental cement. A stainless steel straight headpost was then cemented to the skull posterior to the imaging window.

### Electrode implant

During the window implant surgery, a single-shank 4-channel linear silicon probe (Q-trode, Neuronexus, MI) was implanted to monitor hippocampal local field potentials contralateral to the imaging window. All probes were prepared and plasma sterilized prior to surgery. A 0.7 mm burr hole was first drilled at the coordinates 2.3 mm AP, 2.75 mm ML relative to Bregma, and 0.97 mm DV relative to the cortical surface. The probe was inserted through this hole at a 45° angle lateral to midline, and then cemented to the skull. A stainless-steel jewelers screw served as a ground electrode and was inserted into the right frontal bone of the skull. The connector of the probe was cemented to the headpost at a ~30° angle posterior so as to provide space for the two-photon microscope objective lens.

### Behavioral paradigm

After recovery from surgery mice were water deprived (>85% pre-deprivation weight) and habituated to handling. Next, water deprived mice were head-fixed and trained to run and operantly lick to receive water reward on a cue-less 2m burlap belt. As the animals learned to run on the belt and lick for reward over the course of about 8 days the number of random reward zones was gradually decreased until they reliably ran at least 15 laps in a 15 minute period in a 3 reward zone condition (~8 days). Subsequently, the animals were progressed to a single fixed reward-location on a cue-rich belt (belt Z) different from the belts B and C used for the main experiment (‘goal’ task, see ^13^). For all goal task experiments the rewarded zone was chosen to lie in a cue-poor region of the belts to avoid direct cue-reward pairings. Finally, in order to acclimatize the animal to the full imaging and behavioral set-up the animals were imaged/recorded under the two-photon microscope in the full behavioral paradigm on belt Z for at least 1 day prior to beginning the experimental phase. The full paradigm consisted of three consecutive imaging/recording/behavioral sessions, each 15-20 minutes long. The PRE and POST epochs were conducted on a cue-less burlap belt (belt A) and during these sessions the lickport which was normally used to measure the animal’s licking and deliver reward was retracted, and as a result the animals tended to rest passively rather than actively run. During the intervening RUN epoch animals ran the on a cue-rich belt (belts B or C) for a hidden spatially restricted (one 20 cm zone) operantly delivered water reward. Belts B and C were each made up 3 unique fabrics (6 total fabrics) with unique tactile cues placed along the belts as previously described ^13^. In addition, the belt B and C contexts were further distinguished by the presentation of a unique constant odor during RUN sessions on each of these belts (isoamyl acetate or ‘banana’ odor for belt B and carvone or ‘spearmint’ odor for Belt C). Between the PRE, RUN and POST epochs the animals were briefly (< 5 minutes) removed from head-fixation and placed in their home cages in order to change the belts. One sequential PRE-RUN-POST series was conducted per day. The animals were RUN on belt B for days 1, 2, 3, 8 and 11, and belt C on days 4, 5, 6, 9 and 12 of the paradigm. While the animals were pre-trained in the ‘goal’ task behavioral paradigm, belts B and C were novel to the animal on days 1 and 4, respectively. Offline immobility epochs were defined as those in which the animal’s velocity, smoothed with a half-second Gaussian kernel, was below 3 cm/s for at least 3 consecutive seconds. Online running epochs were defined as those in which the animal’s smoothed velocity was above 5 cm/s for at least 3 consecutive seconds.

### LFP recording and processing

All electrophysiological recordings were performed at 20 kHz, using a multiplexed digital acquisition system (Intan RHD). Only mice showing clear LFP-recorded SWR’s events were included in this study. Electrophysiological and LFP recordings were synced to the calcium imaging by recording the imaging system’s ‘Pockels cell’ signal occurring at the end of each calcium imaging frame. LFP signals were calculated from the wide-band 20 kHz signal by down-sampling to 1250 Hz. From the 4 sites on the silicon probe the pyramidal layer site was identified as that showing the greatest ripple-band Gabor wavelet power (100 to 225 Hz). After the removal of noisy LFP epochs, SWR events were detected using a costume Matlab supervised algorithm based on a template hand-labeled ground-truth data set of SWR events from 4 mice (separate from those recorded in this study) using k-nearest-neighbor embedding based on wavelet-derived SWR features. Candidate events identified using the supervised template matching procedure, were considered SWR-events if their within-event ripple-band wavelet power was at least 12 median absolute deviations (m.a.d.) above the median for the session. SWR-event detection was visually verified. Only SWR-events occurring during periods of immobility lasting at least 3 seconds were included in the analysis. For calculations requiring point time estimates, such as SWR-triggered peri-event time histograms (PETH’s), the within-SWR ripple-power peak was used. In order to account for SWR-bursts in which long-reactivations may occur ^19^ SWR events occurring within 200 ms of each other were merged. For theta phase estimation, wavelets occurring during RUN running-bouts were taken between 5 and 10 Hz at 0.5 Hz intervals. The instantaneous theta phase at time *t* was estimated from the imaginary wavelet component for the frequency showing maximal absolute wavelet power at time *t*. This procedure resulted in a flat distribution of observed theta phases over time (data not shown).

### Calcium imaging and data processing

All imaging was conducted using a two-photon resonant scanner (Bruker) using. Calcium imaging was acquired at 60 Hz (512 × 256 lines per frame). The excitation laser was 920 nm (50-100 mW, Coherent Ultra II). A preamp (1.4 × 105 dB, Bruker) was used to amplify signals before digitization. Pockels cells were used to regulate the power of the laser reaching the tissue. Images were acquired using a Nikon 40X NIR water-immersion objective (0.8 NA, 3.5 mm WD) at 1-1.2 zoom, resulting in FOV’s of 320×320 µm to 267×267 µm. At the beginning of each imaging session the animal’s FOV was carefully relocated and centered by comparison to a reference image. Each imaging session was motion corrected by maximizing each whole-frame’s cross-correlation to the average frame. In order to assess the quality of motion correction and exclude frames containing z-motion, the correlation coefficients of individual frames to the average motion corrected frame was calculated and low correlation frames were excluded (2.5% of frames) – these epochs were also censored in subsequent analyses. Next, the three sessions recorded each day were aligned using a cross-correlation maximizing 2-dimensional affine transform and concatenated with only pixels observed across all imaging sessions for a given day used for further analysis. The concatenated movies were then processed using the Suite2p software package ^33^ to identify active spatial masks corresponding to neural regions of interests (ROIs) and extract associated fluorescence signals. Identified ROIs were curated post-hoc using the Suite2p graphical interface.

For cross-day ROI registration, the average frame was calculated for each whole-day concatenated movie, and all cross-day pair-wise affine transforms and resulting average frame correlation coefficients were taken. The day with the highest correlation to other days and the lowest percentage of non-overlapping pixels was chosen as the reference, and affine transforms (calculated from the per-day average frames) were applied to the ROIs such that they were all 20 aligned to the reference day. Subsequently, a modified version of the analysis laid out in ^34^ was applied to determine groups of cross-day ROI-pairs putatively corresponding to the same physical cells. Briefly, each ROI was binarized as the minimum subset of pixels accounting for at least 95% of the total ROI mask weight. Candidate cross-day binarized ROI pairs were identified as those whose distance as measured by their closest pair-wise pixels was not greater than 12µm ^34^. The Jaccard distance scores for all candidate ROI pairs were calculated. For each animal, the bi-modal distribution of a Jaccard distances were jointly fit by as Beta and Gamma distributions, with the Beta distribution modeling high Jaccard distance scores corresponding to ROI-pairs putatively belonging distinct cells and the Gamma distribution modeling low Jaccard distance scores corresponding to ROI-pairs putatively belonging to the same cell imaged across days. For each pair, the Bayesian posterior probability of belonging either to the Beta or Gamma distribution was calculated as described in ^34^. These pair-wise posterior probability values were hierarchically clustered into groups of ROIs putatively corresponding to the same cells using an inter-cluster average-distance cutoff of 0.99 posterior probability of belonging to the Gamma (Same-Cell) distribution. The resulting clusters were extensively verified by visual inspection. Following the Suite2p based ROI detection and the ROI cross-registration, each day’s imaging sessions were de-concatenated and further analysis was performed on a per-session basis.

### Calcium activity detection

In order to reduce calcium white ‘scatter’ noise and improve signal detection, a wavelet-based denoising algorithm (MATLAB 2019, *wden* function, using parameters, ‘*modwtsqtwolog’,’s’, ‘mln’, 5, ‘sym4’*) was applied to the calcium activity traces resulting in the denoised activity trace vector *T*_*dn*_. This algorithm was adopted as, when compared to other algorithms such as Savitsky-Golay filtering, it resulted in robust white noise reduction while minimally altering the underlying waveforms of observable calcium transient events. Subsequently, the per-trace median was subtracted from each of the smoothed traces and they were deconvolved using a first-order auto-regressive model as implemented in the OASIS software package ^35^. This algorithm outputs the deconvolved (spike estimate) of the trace *C*_*sp*_, and a denoised trace reconstruction *T*_*est*_ based on the re-convolution of the spike estimates. The deconvolution noise was taken as the median absolute deviation (mad) of the residual of the observed trace *T* and the reconstructed trace *T*_*est*_. Spike estimates, *C*_*sp*_, were normalized by the deconvolution noise and binarized by thresholding, resulting in the binary spike estimate vector *S*_*bi*_. Based on the observed differences in calcium activity waveforms between the online and offline epochs (see fig. S2), a threshold of 1.5 mads was used for online running epochs, while a lower threshold of 1.25 mads were used for offline immobility epochs. Since small inter or intra-cell variability in transient amplitude and duration can lead to substantially different numbers of thresholded estimated spikes, a further sparsification step was applied in which for any consecutive frames in which thresholded spikes were detected only the estimated thresholded spike corresponding to the first frame of the sequence was kept, resulting in the sparsified binary spike estimate vector *S*_*sp*_ which, unless otherwise stated, is the calcium-based activity estimate used in this study.

A separate transient-based analysis was carried out as a control for the spike estimate-based analysis. Briefly, since large amplitude transients can bias baseline estimation, for each denoised trace vectors, *T*_*dn*_, large amplitude transients with peaks 15 mads above the median (edges at 4 mads above the median) were detected. The large amplitude transients were temporarily removed and the median and mad values were recomputed from these censored traces. Online running related transients were detected as those with peak amplitudes of 14 mads above the censored medians, while offline immobility related transients were detected as those with peak amplitudes 6 mads above the censored medians with the edges of both set at 1 mad above the censored medians. Finally, since sequential transients may occur before a given ROI returns to baseline fluorescence levels, a final splitting step was performed in which local minima in transients were detected as points with local trough prominences (calculated as the peak prominence of the inverted within-transient traces) exceeding 0.2. The mad-based threshold criteria outlined above was reapplied to the split transients. Transient peak times were used as point estimates of ROI calcium transient based activity.

### Place field detection and stability measurements

The animal’s position during online running epochs on the 2m RUN belts was binned into 100, 2 cm spatial-bins. For each cell, as within spatial-bin firing rate was calculated across all bins based on its sparsified spike estimate vector, *S*_*sp*_. This firing-rate by position vector was subsequently smoothed with a 7.5 cm Gaussian kernel leading to the smoothed firing rate by position vector. In addition, for each cell, 2,000 shuffled smoothed-firing rate by position vectors were computed for each cell following the per-lap randomized circular permutation of estimated activity vector, *S*_*sp*_. For each cell, putative place fields (PFs) were defined as those in which the observed smoothed firing rate by position vectors exceeded the 99^th^ percentile of their shuffled smoothed firing rate by position vectors as assessed on a per spatial-bin basis, for at least 5 consecutive spatial bins. As an additional control, only those putative PFs in which the cell had a greater within-PF than outside-of-PF firing rates in at least 3 or 15% of laps (whichever was greater for each session) were considered *bona fide* PFs and kept for further analysis. Cell’s exhibiting at least one PF were considered place cells (PCs). PC’s spatial information per second and per estimated spike was calculated as described previously ^36^ both for the observed and shuffled firing-rate by position vectors. Since this measure can be strongly skewed in the low firing rate regime observed in calcium imaging, in the extreme case leading to high information scores for cells which only fired once ^32^, for each cell the observed information scores were normalized as the z-score relative to information scores observed in their shuffled data. Place coding was not assessed for PRE or POST epoch sessions as the animals were not rewarded during these epochs and thus tended not run. Unless otherwise indicated same-cell pair cross-day spatial firing stability for *i*^*th*^ cell imaged on days *N* and days *N* + *B* was calculated as the spatial distance in centimeters between the cell’s highest-firing rate within PF peaks on day *N* and day *N* + *B*. In order to account for spurious elevated stability resulting from non-uniform PF coverage of the belt, this distance was z-score normalized by (mean subtracted and divided by the S.D. of) the PF peak distances between *i*^*th*^ cell’s PF peak distance with all other place cells imaged on both days *N* and days *N* + *B*. Finally, the sign of the resulting normalized stability score was flipped such that larger values corresponded to more highly preserved spatial coding, resulting in the stability units of ‘z-scored – Δcm’.

Place field onset laps (fig. S4, E-I) were determined as in ^37^. Briefly, for each place cell the per lap within-PF to outside-of-PF firing rate difference was calculated. The PF onset lap was defined as the first lap showing higher within than outside of PF firing rate of the first sequence of 6 laps in which the within-PF firing rate was higher than the outside-of-PF firing rate for at least 4 of the 6 laps.

### Bayesian Reconstruction Analysis

Bayesian reconstruction of virtual position ^19^ was performed utilizing a template comprising of all place cell’s smoothed firing rate-by-position vectors as:

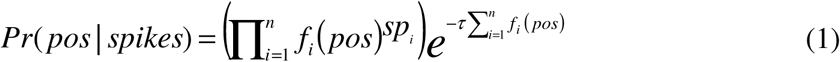

Where *fi(pos)* is the value of the firing rate-by-position vector of the *i*^*th*^ place cell at position *pos*, *sp*^*i*^ is the number of spikes fired by the *i*^*th*^ place cell in the time bin being decoded, τ is the duration of the time bin and *n* is the total number of place cells. Time bins (τ) of 20 frames (~333 ms) were used for Bayesian reconstruction of online activity, while a time bins of 2 frames (~33 ms) were used for Bayesian reconstruction of offline activity and only bins with non-zero firing rates were used for offline Bayesian decoding. Posterior probabilities were subsequently normalized to one:

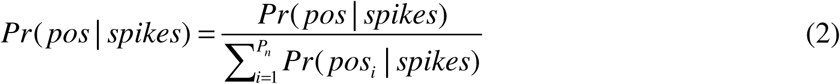

Where *P*_*n*_ is the total number of positions (2 cm bins). When assessing reconstruction accuracy, or reconstructed distance to the reward, the position corresponding to the maximal posterior probability was taken. For same-day decoding a 5-fold (applied by lap-number) cross-validation was used. For cross-day position reconstruction only registered cells which were found to be place cells on both the training and testing days were included, and only pairs of sessions containing at least 5 such place cells were included in the decoding analysis.

### Pair-Wise and ICA Ensemble Reactivation

For pair-wise reactivation analysis the RUN PF peak distance between pairs of place cells was compared to their offline firing-rate Pearson’s correlation coefficents in either the PRE or POST epochs. For calculating offline firing rate correlations the sparsified binary spike estimate vectors *S*_*sp*_ were restricted to periods of immobility during either PRE or POST and convolved with a 150 ms Gaussian kernel. ICA ensembles were detected using previously described methods ^17,18^. Briefly, place cell RUN running-bout spike estimate vectors, *S*_*sp*_, were convolved with a 1 second Gaussian kernel corresponding to behavioral time-scales. Subsequently, the ICA components of these smoothed RUN activity estimates were calculated across time using the fastICA algorithm ^38^. Only the subset of components found to be significant under the Marcenko-Pasteur distribution were considered ICA ensembles and included in the ICA ensemble matrix, *w*, with rows corresponding to place cells, and columns corresponding to significant ICA components. For each component, *b*, of ICA ensemble matrix *w* a square projection matrix, *P*, was computed from *w*_*b*_ as:

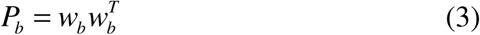

Where *T* denotes the transpose operator. Subsequently, the diagonal of the projection matrix *P* was set to zero to exclude each cell’s individual firing rate variance. Offline reactivation was assessed from the 150 ms Gaussian kernel convolved offline activity matrix *Z*. For the *i*^*th*^ time point (frame) in *Z* the reactivation strength *R*_*b,i*_ of the *b*^*th*^ ICA component was calculated as the square of the projection length of *Z*_*i*_ on *P*_*b*_:

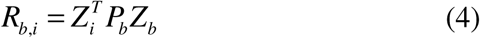

In order to assess the *x*^*th*^ cell’s contribution to ICA reactivation, a per cell contribution (PCC) score was defined as the mean across all components *b*, and time-points, *i*, of the reactivation score, *R*, computed from all place cells, *c*, minus the reactivation score *R*_*c\x*_ computed after the exclusion *x*^*th*^ cell from the activity and template matrices. To account for putatively non-specific changes in ICA reactivation strength from the PRE to the POST epochs, normalized PCC was taken per session as the PRE to POST change in within-epoch PCC rank. ICA components were shuffled by randomly permuting the weight matrix *w* across place cells and recalculating the reactivation strength. When ICA ensemble analysis was applied across days *N* and *N* + *B*, day *N* ICA ensemble matrices *w* were recomputed based on the subset of day_*N*_ place cells which were also registered on day_*N*+*B*_.

### Population synchrony events (PSE’s) and offline sequence detection

Offline PSE’s were detected by convolving each place cell’s (as assessed during that day’s RUN) offline immobility firing rate vector *S*_*sp*_ with a 125 ms Gaussian kernel and z-scoring the smoothed firing rate vector. Subsequently, for each frame *i* the population mean of the smoothed and z-scored vector was taken across place cells, and subsequently z-scored. Putative PSE’s were defined as epochs during which the z-scored population activity vector reached a peak of at least 3.5 SD’s. above the mean with event-edges at 1 SD above the mean, with a minimum inter-event time of 0.2 seconds. Only PSE events lasting between 0.2 seconds (12 frames) and 1 second (60 frames), and during which at least 5 distinct place cells each fired at least one estimated spike, were kept for further analysis.

In order to perform offline sequence analysis within-PSE place cell activity was binned into non-overlapping 2-frame time-bins, and Bayesian decoding was performed on these two frame bins as described above. Sequence content is commonly measured as the weighted linear correlation, weighted_r(linear)_*(pos, bin*; *Pr*), between time and decoded position as weighted by the posterior probability of position ^21^. For a given PSE the weighted mean *m* is given by:

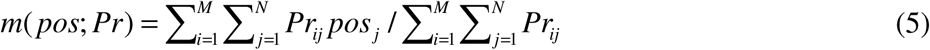

And the weighted covariance *cov*:

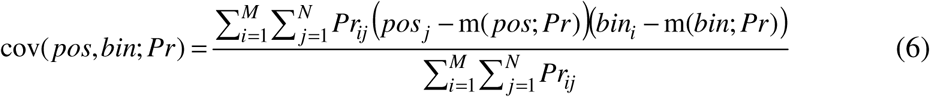

And finally the weighted linear correlation weighted_r(linear)_*(pos, bin*; *Pr):*

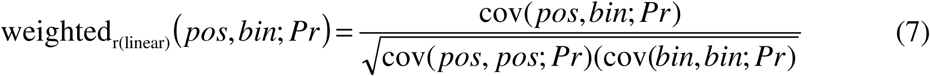

Where *pos*_*j*_ is the *j*^*th*^ spatial bin, *bin*_*i*_ is the *i*^*th*^ temporal (2 frame) bin in the event, *Pr*_*ij*_ is the Bayesian posterior probability for that spatial bin at that temporal bin, *M* is the total number of temporal bins and *N* is the total number of spatial bins. However, this measurement only accounts for linear relationships and is thus not sufficient to detect sequences of the circular RUN belt which may span the artificially defined belt ‘edges’. Generally, the circo-linear correlation coefficient, *r*_*cl*_, between a circular variable *α*, and a linear variable *x*, is defined as:

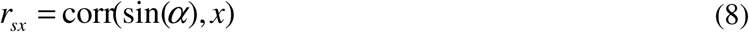

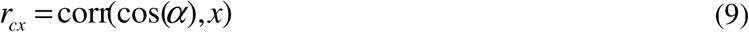

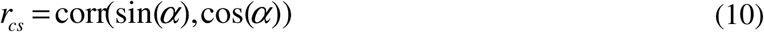

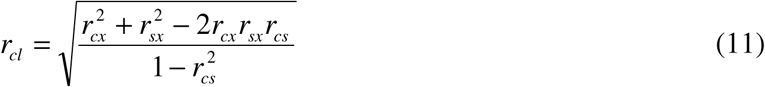

Where ‘corr’, ‘sin’, and ‘cos’ denote the Pearson’s (linear) correlation, sine and cosine operators, respectively. Therefore, the circo-linear weighted correlation coefficient between time (the linear variable) and position (the circular variable) weighted by the posterior probability of position in each time bin was derived by combining equations 5–7 and 8–11 as:

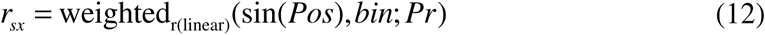

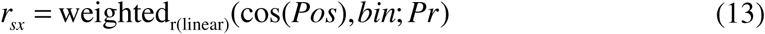

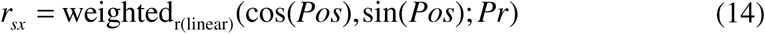

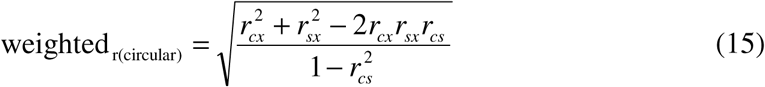

For each event, the observed weighted circo-linear correlation coefficents weighted_r(circular)_, hereafter referred to as weighted-r, was compared to a distribution of 2,000 null weighted-r values either by z-scoring the observed value by the null values (rZ score) or as the empirical p-value. While several shuffle approaches were used (see fig. S11 legend) the principal shuffle used in the main figures involved the random re-ordering (resampling without replacement) of the bins observed within a given event. In order to determine the precise trajectory content of each sequence, a modified ‘line-casting’ or Radon transformation ^19^ approach was employed. Briefly, for each event the Posterior probabilities were tiled twice by position to account for ‘edge’ spanning sequences, and smoothed with 5 cm Gaussian kernel across positions within each time-bin. Subsequently, lines, restricted to those crossing all bins, were densely cast along this matrix and the mean of the Posterior probability for each of these lines was calculated. The trajectory was defined as the casted line with the highest mean posterior probability value and the sign of the slope of the trajectory line defined whether it was a forward or reverse sequence.

### SWR recruitment and future stability analysis

In order to examine the interaction between SWR recruitment and future stability, two complementary methods for selecting groups of Low and High ΔSWR-gain cells were utilized. In the ‘per-cell’ selection method for each reference RUN session cells were restricted to those meeting the following criteria: 1) the cell was place cell on that session, 2) the place cell was novel, that is, the place cell had never previously shown a significant place field on previous sessions on the same belt (either B or C), 3) the place cell showed significant spatial coding (i.e. was a place cell) in at least one future RUN session on the same belt. Furthermore, since the objective of the analysis was to assess the interaction of the PRE to POST changes in SWR recruitment and future place coding stability, 4) place cells were restricted to those which had none-zero within-SWR firing rates across the entire imaging day (84.6% of the cells meeting criteria 1-3 above). For each session, place cells meeting criteria 1 through 4 above were split by their POST – PRE change in within-SWR gain (their within-SWR firing rate in either PRE or POST divided by their overall immobility firing rate in either PRE or POST, respectively) median such that the Low and High Δwithin-SWR gain groups had an equal number of place cells from each session. This balanced designed controls for putatively non-specific variance in future spatial coding stability accounted for by individual sessions or animals. For each of these cells future place coding stability was assessed as the mean of the coding stability between its place coding on the reference day RUN’s session to future days the same cell was a place cell on the same RUN belt. For the Goal and Path zone analyses, place cells were first divided into three groups: Reward zone cells (≤ 10 cm PF peak reward distance, not used for further analysis), Goal zone cells (between > 10 cm and ≤ 55 cm PF peak reward distance) and Path zone cells (> 55 cm PF peak reward distance). This corresponds to the splitting of the belt into two equal-length (each 90 cm total length) portions based on proximity to the reward, after the exclusion of the 20 cm Reward zone. The ‘per-cell’ analysis described above was carried out independently on the Goal and Path zone cells. The significance of the interaction between the Path/Goal zone cell group, Low/High ∆SWR-gain cell group and a future place field stability was computed via permutation test in which the observed difference in Path cell [High – Low future stability] versus Goal cell [High – Low future stability] was compared to the same value computed in 25,000 permuted data sets in which High vs. Low ∆SWR-gain cell labels were randomly permuted as a one-sided empirical p-value.

While the ‘per-cell’ approach above insured that each cell contributed to the analysis at most twice (once for each RUN belt) a complementary ‘per-same-cell-pair’ approach was included that instead compared each pair of days that given cell was a place cell on the same belt independently. Briefly, for each pair of days *N* and *O* (where *N* < *O*) on which the animals ran on the same RUN belt (B or C) cells were included in the analysis if they 1) were place cells on both days *N* and *O*, and 2) fired in at least one SWR event on day *N*. For cells meeting criteria 1 and 2 above, cells were split by the median their POST – PRE change in within-SWR gain as described above. This procedure was repeated for all day *N* and *O* pairs which for which the animals ran on the same RUN belt. As in the ‘per-cell’ approach above, the ‘per-same-cell-pair’ approach balanced design controls for day *N* inter-session variability, while also additionally balancing day *O* inter-session variability.

### Generalized linear model (GLM) predicting stability

In order to test the independent contribution of PRE to POST changes in SWR-recruitment and other factors which may be relevant to future place coding stability a generalized linear model predicting future place field stability was built. Briefly, for the subset of novel place cells described in the ‘per-cell’ analysis above, a Ridge regression-based GLM was built with the following factors: 1) the cell’s spatial information per second on the novel place field day, 2) the mean days apart between the novel place field day and the future days on the same RUN belt with which stability was compared, 3) the cell’s normalized within session stability (z-scored – Δcm between the first and second half of laps on the novel place field day), 4) the cell’s PF peak distance to reward on the novel place field day, and 5) the PRE to POST change in within-SWR gain observed on the novel place field day. In addition, the interactions between the 5 factors were also used as regressors, for a total of 15 regressors. While additional factors were tested (data not shown), these 5 factors were chosen for clarity as representing a wide range of potentially independent determinants of mutli-day coding stability. Individual regressors were linearized and scaled by ranking followed by z-scoring within each session. Ten-fold cross-validated Ridge-regression GLM’s predicting linearized and scaled future place coding were constructed based on the 15 regressors. Subsequently, the contribution of each regressor was assessed by recomputing the 10-fold cross-validated GLM after the exclusion of each regressor individually. The significance of the reduce versus full models was tested by two complementary methods: 1) an F-test was run based on the difference of mean squared errors in the full versus reduced models, 2) full versus reduced model differences in median absolute error were compared on a per-fold basis.

### Data Availability Statement

Processed imaging and LFP data and analysis code for this study will be made available through the Neurodata Without Borders (www.nwb.org) service.

## Acknowledgments

We thank G. Buzsáki, J. Magee, D. Levenstein, J. Priestley, K. Kay, and S. Tuncdemir for invaluable comments and discussions on the analysis and the manuscript. We thank J. Bowler for the behavioral software used in these experiments. This work was supported by the Brain Initiative Grant NIH U19NS104590 and by the Charles H. Revson Senior Fellowship in Biomedical Science. Authors declare no competing conflicts of interest.

## Author Contributions

A.D.G and A.L. designed the experiments and prepared the manuscript. A.D.G. performed the computational analysis. A.D.G., F.T.S. and M.J.D performed the experiments.

