## Supplementary Information for "Offline Memory Reactivation Promotes the Consolidation Of Spatially Unbiased Long-Term Cognitive Maps"

**This PDF file includes:**

Figs. S1 to S16

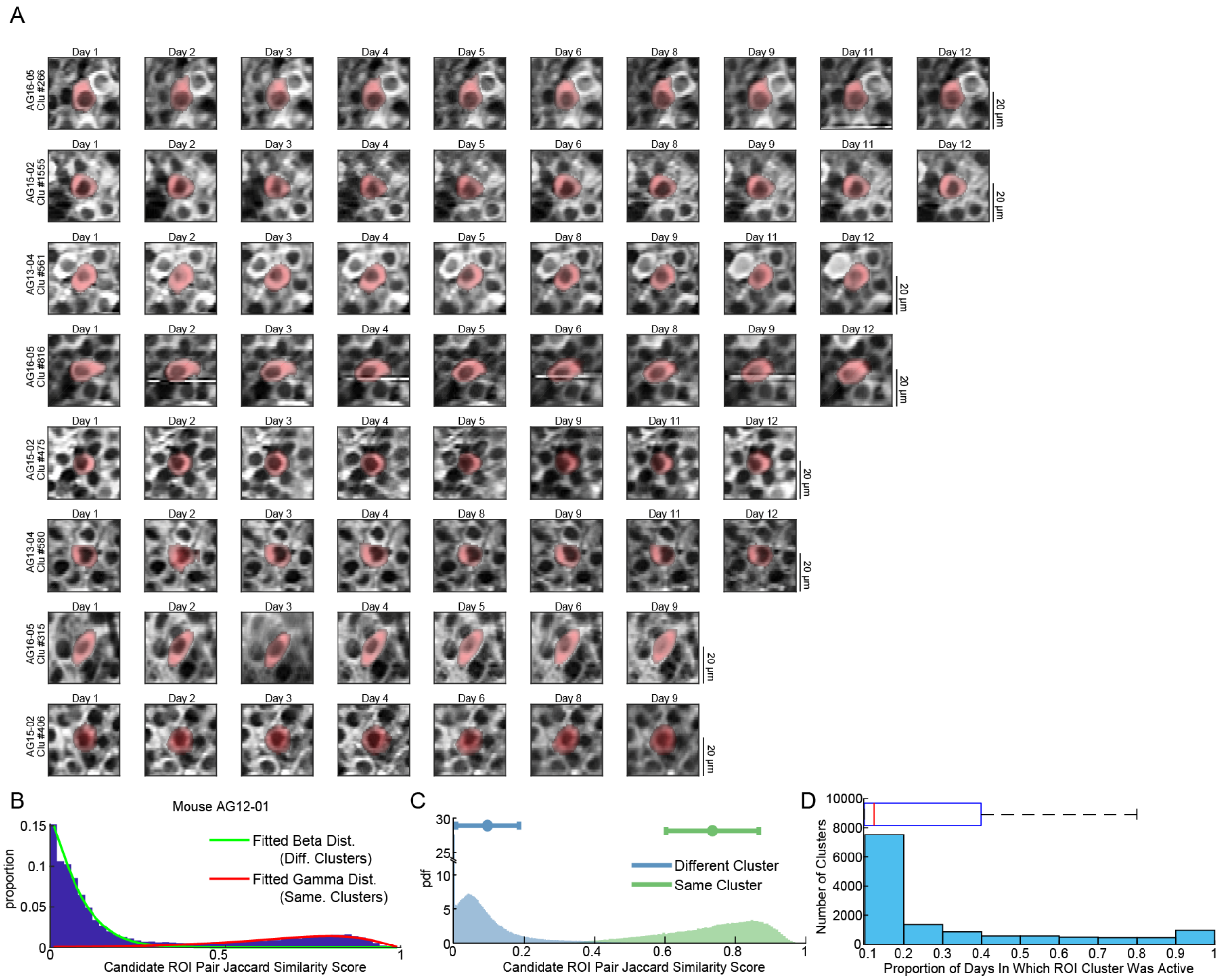


**Fig. S1: Cross-day ROI registration.** **(A)** Eight examples of same-cell registered ROI's imaged over 9, 8, 7, and 6 days (top 2 rows, second 2 rows, third 2 rows, fourth 2 rows, respectively). **(B)** Cells were registered using an adapted version of the algorithm introduced in [1]. Briefly, a beta and a gamma distribution were jointly fit to the Jaccard similarity scores (number of overlapping pixels, divided by the total overlapping and non-overlapping pixels) of cross-day ROI pairs located ≤12µm away from each other (as measured from their nearest pixels). The panel shows the beta and gamma distribution fits for one animal, showing that the joint distribution approach well matches the bi-modality of observed in the scores. The posterior probability of belonging to either of these distributions was subsequently used to cluster ROIs into cross-day groups putatively belonging the same cell imaged over multiple days. **(C)** Group data for pair-wise Jaccard similarity scores for pairs of ROI's belonging to different cell-clusters (blue, mean: 0.09, median: 0.06, n=302,350 ROI-pairs, horizontal bars show mean±SEM) or to the same cell-clusters (green, mean: 0.73, median: 0.76, n=101,222 ROI pairs). Note the clear separation between same-cell clustered and non-clustered Jaccard similarity scores. **(D)** The histogram shows the distribution of the proportion of imaging days each ROI-cluster was imaged for (horizontal overlay shows the box-and whisker plot, red-line shows median, blue box shows $25^{th}$ and $75^{th}$ percentiles, dashed line shows the $90^{th}$ percentile). While due to the conservative approach to clustering (panels B and C) 41.8% of putative pyramidal cells were observed in only one imaging day (i.e. were not clustered with other ROI's), 32.3% of putative pyramidal cells were observed across at least 1/3rd of imaging days, and 22.2% putative pyramidal cells were observed in at least half of the imaging days providing a substantial sample in which to examine cross-day pyramidal cell dynamics.

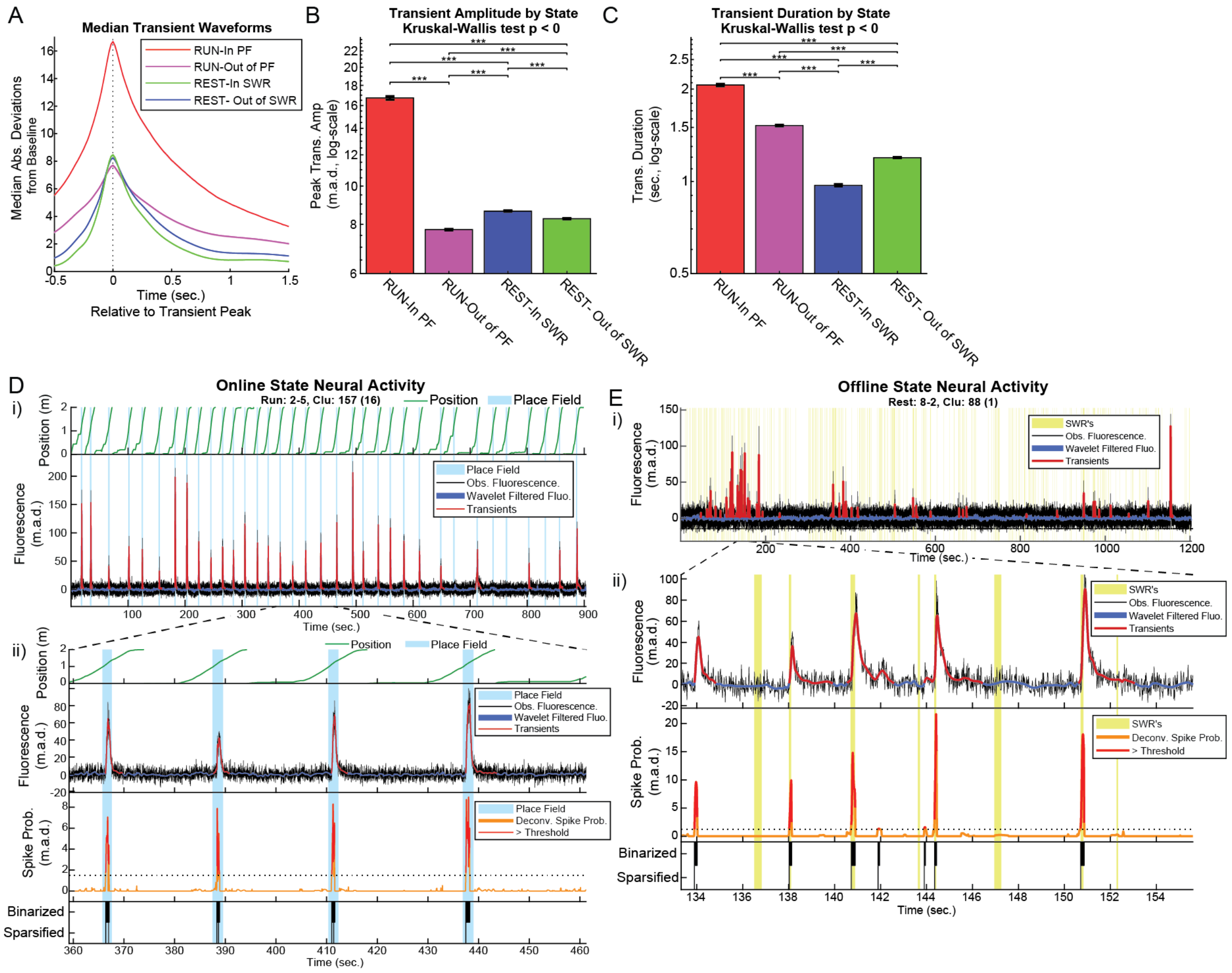


**Fig. S2: Hippocampal calcium Activity varies by behavioral state.** In order to conduct a comparison of calcium neural responses in different behavioral states, preliminary transients were detected using a 5 m.a.d. (median absolute deviation) threshold of 5. **(A)** Marked differences were observed in the median transients (shown ± boot-strapped 95% confidence interval) between different states (n = 319,846 total transients, the behavioral states of transients were determined by the time-point of the transient peak). (**B**) Notably, within place field (PF) transients during the running were found to have the largest amplitude, followed by immobility transients occurring within LFP-detected SWR-events, with outside of PF transients occurring during running, and non-SWR-related transients occurring during immobility displaying significantly lower amplitudes (Kruskal-Wallis test, followed by post-hock Tukey-Kramer tests). **(C)** Likewise, within PF running transients showed the longest duration, however, SWR-related transients were of the shortest duration when compared to other states, likely reflecting the short-duration of SWR-related excitability burst as well as the relative sparsity of pyramidal cell bursting during SWR's [2]. Due to the observed differences in transient amplitude and duration between states, as well as the decreased potential for motion related contamination during immobility epochs, a lower threshold for activity was used for offline (immobility) as compared to online (running) related epochs. **(D)** Example of activity estimation for a place cell during an online RUN epoch. (D, i) The top panel shows the position of the animal on the belt during the RUN session (green lines), with epochs during which the animal was running within the cell's place field shaded in blue. The bottom of panel D-i shows the m.a.d.-normalized fluorescence trace of the cell (black) as well as the wavelet denoised trace used for activity estimation. Significant online transients (m.a.d. ≥14) are highlighted in red. The top two panels of D-ii show a zoomed in portion of the panels shown in (i). The third panel in (ii) shows the deconvolved spiking probability (orange), with supra-threshold spike probability events (m.a.d. ≥1.5) shown in red. The bottom panel in (i) shows the binarized (top-row) spike rasters, as well as the final sparsified binarized raster which was used as the estimate of cellular activity in the current study, unless otherwise specified. **(E)** Example of activity estimation for a cell during an offline POST epoch. Panel (E, i) shows the whole POST session fluorescence trace (black), the wavelet denoised trace (blue) as well as the detected offline transients (m.a.d. ≥6) – SWR events are shaded in yellow. Panel (E, ii) shows a zoomed in portion of the POST session. Note that transient activity is enriched within, but is not limited to, LFP-detected SWR events. A threshold of 1.25 m.a.d. was used to detect deconvolved spiking events during offline epochs, and these binarized events were subsequently sparsified as described in the Online Methods.

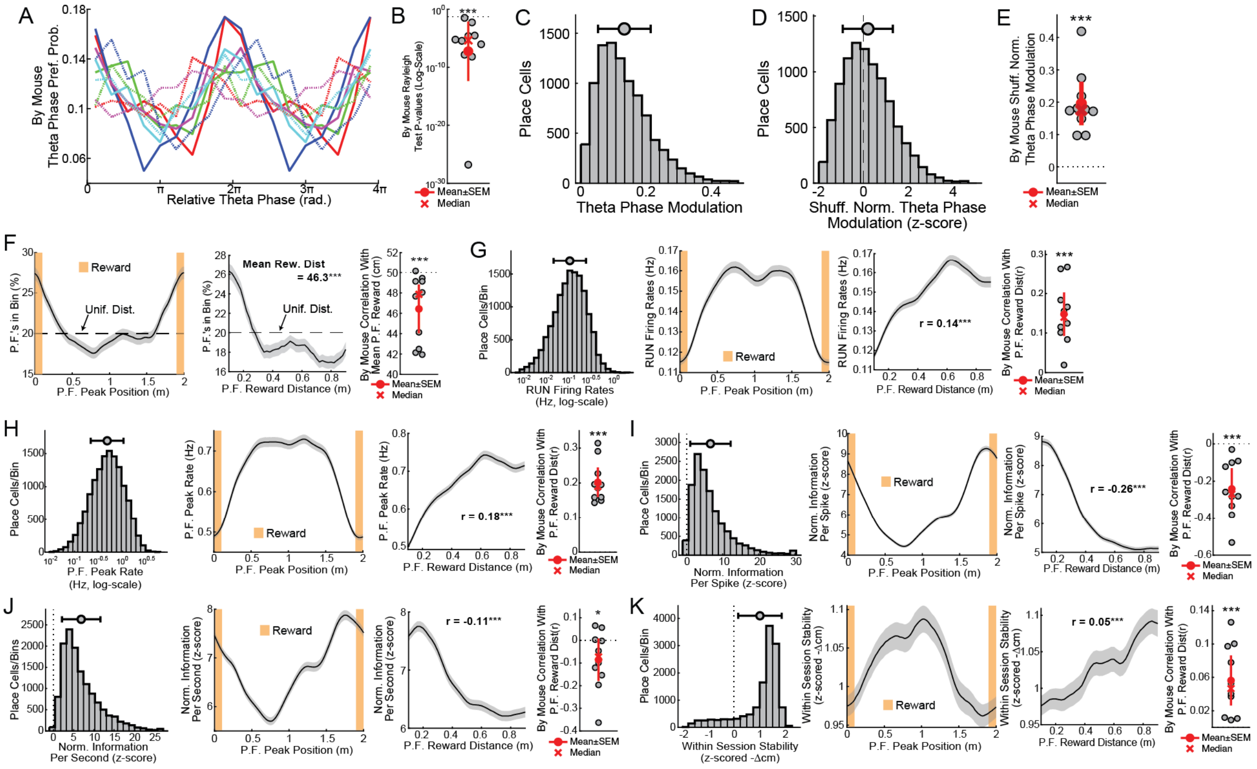


**Fig. S3. Place cell online calcium activity is modulated by the theta oscillation and by distance to reward. (A)** Distribution of place cell preferred normalized theta phases during online running (see Fig. 1F), each line shows an individual animal (n = 10 mice). **(B)** Per animal p-values of Rayleigh tests for uniformity of a circular distribution on the per cell theta phase preferences (group p-value ~ 0 is the one-sided boot-strapped test of the difference of the mean of the per animal p-values from 0.05, n = 10 mice). **(C)** Distribution of theta phase modulation amplitudes (resultant circular vector lengths, mean: 0.20, median: 0.17, horizontal bars show mean ±SEM). **(D)** For each cell the observed theta modulation amplitudes were compared to a null distribution of 1,000 theta modulations amplitudes obtained from a circular shuffling of the included epochs. For each cell the observed modulation values were z-scored relative to their null distributions such that the expected normalized modulation due to chance was zero (observed mean: 0.19, median: 0.07, Signed-Rank test p~0). **(E)** The mean normalized modulation score per animal confirm that theta modulation of the calcium was consistent throughout our dataset (group p-value ~ 0 is the one-sided boot-strapped difference of the per animal means from the expected null of 0, n = 10 mice). **(F,** left panel**)** The distribution of place field (PF) peak coding locations is plotted as the percentage of place cells with PF peaks within a circularly sliding ±20 cm window (percentage ±boot-strapped 95% confidence interval). Note the enrichment of PF's near the rewarded location (dashed line shows the expected percentages based on a uniform distribution of PF's along the entire belt). (F, middle panel) This enrichment was confirmed by plotting the percentage of place fields found at difference distances from the reward location (percentage within a 20cm sliding window ±boot-strapped 95% confidence interval, p-value is the Signed-Rank test of the observed PF distances to reward from the 50cm distance observed under uniform PF tiling of the belt, n = 13,341 place cells). (F, right panel) For each animal the mean PF distance from reward is plotted, confirming the enrichment of PF's near the reward (p-value ~ 0, is the one-sided boot-strapped difference of the per animal PF reward distance compared to the 50cm distance expected under a uniform distribution, n = 10 mice). **(G**, first panel**)** The distribution of place cell firing rates during running bouts during the RUN epoch (mean: 0.07 Hz, median: 0.039 Hz, n = 13,341 place cells). (G, second panel) The mean online firing rate of place cells during running epochs is plotted within a circularly sliding ±20 cm window (mean ±SEM). (G, third panel) The mean online firing rate is plotted at different distances to the reward (20 cm sliding window ±SEM, r-value from Pearson's correlation between online firing rate and PF distance to reward across cells, n = 13,341 place cells). **(H)** The relationship between PF peak firing rate, position and distance to reward is plotted as in G, mean peak firing rate: 0.64 Hz, median: 0.49 Hz). Both overall and peak firing rates showed significant up-regulation farther from the reward zone. **(I)** The shuffle-normalized spatial information per estimated spike of place cells is shown as a function of position and distance to reward (plotted as in panel G, mean normalized information per spike: 6.6, median: 5.1, 0 is the value expected by chance). **(J)** The shuffle normalized spatial information per second of place cells is plotted as a function of position and distance to reward as in panel G (mean: 6.9, median: 5.5). Notably, the spatial information measures shown in panels I and J display the opposite relationship to reward distance as do the firing rate measures shown in G and H. **(K)** Finally, the normalized within session stability (first half of laps compared to second half of laps) is shown as a function of position and PF distance to reward as in panel G (mean: 1.02 a.u., median: 1.35 a.u.) showing a slight but significant positive relationship with distance to reward. Therefore, while reward-distant cells are less numerous and less spatially informative, they have higher firing rates and code slightly more stably across space within individual sessions.

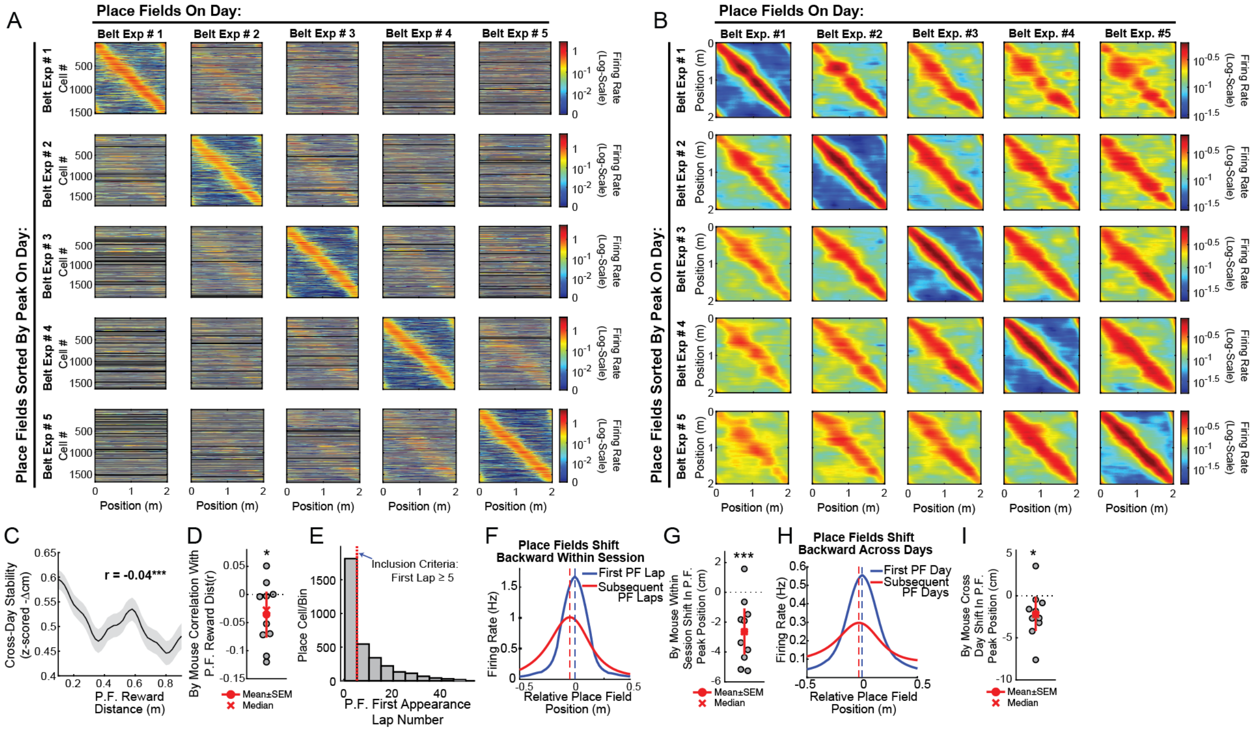


**Fig. S4. Place cell coding dynamics across time shows both stable and unstable elements.** **(A)** The firing rate by position vector of place cells is shown across RUN belt exposure numbers (number of days/sessions on a given RUN belt). The firing-by position vectors observed on the belt exposure number shown on the top of the panel were sorted according to their place field peak firing position on the belt exposure number shown on the left of the panel. Only place cells with significant place coding on at least two RUN sessions on the same belt were included and the data was pooled across both RUN belts. Cells that were not imaged or did not show significant place coding on a given day are colored in black. As the difference between the reference belt exposure number (top labels of panel) and the comparison belt exposure number (left labels of panel) increases, the place fields become less stable (see Fig. 1A). **(B)** The same data in panel (A) is shown after applying a ±10 cm rolling average across reference place field peaks (panel A, subpanel y-ordinate). Applying this smoothing reveals that while the spatial code partially decays across time, a substantial amount of stable place coding persists across days. **(C)** The cross-day stability of place cells on the same belt is shown as a function of place field peak distance to reward on the first day of the comparison, shown within a 20cm sliding window (±boot-strapped 95% confidence). A weak but significant effect was observed such that, unlike within-session stability (fig. S3, K), place cells with peaks further from the reward tended to show decreased cross-day spatial stability (Pearson's correlation between P.F. peak reward distance and cross-day stability (r): -0.04, $p < 9.3\times10^{-5}$, n = 9,721 cross-day pairs). **(D)** This relationship was found to be present across animals. Plots show the correlation coefficents as shown in (C) but within mice (p-value from one-sided boot-strap test of the mean, $p < 0.017$). **(E)** Growing evidence points to behavioral time-scale synaptic plasticity (BTSP) as a central mechanism of place field formation [3]. One of the consequences of this mechanism is that place fields are observed to shift backwards in space on laps subsequent to the place field formation lap. In order to examine this phenomenon we calculated the place field onset lap (see Online Methods) for novel place cells, that is for days in which a given place cell displayed its first place field for a given RUN belt. In order to examine *de novo* PF formations events on the belt, only novel place cells showing PF formation events at or after the fifth lap of the RUN were included for further analysis (mean PF formation lap: 8.58, median: 5, n = 3,316 novel place cells). **(F)** Place cell firing rate by position vectors are shown relative to the peak firing position on the first lap (graphs show mean±SEM). It was found that, consistent with a BTSP-like mechanism, place cell coding location tended to shift backwards (negative values) on laps subsequent to the formation lap (mean shift: -3.3 cm, median shift: -4 cm, Signed-Ranked test p ~ 0, n = 1,672 *de novo* place cells). **(G)** The backward shift from the formation lap to subsequent laps was found to be consistent across animals (one-way boot-strap test of the mean, $p < 4x10^{-5}$). **(H)** However, since it is not clear whether similar mechanisms govern cross-day spatial coding we calculated place cell's firing by position vectors either on the first day that the place cell showed a place field on a given RUN belt or for the mean of subsequent days in which the same place cell showed place fields on the same belt relative to the PF peak firing location on the first PF day. Notably, a similar backward shift was observed (mean shit: -2.3 cm, median shift: -2.41 cm, $p < 9.4x10^{-5}$, n = 2,916 novel place cells). **(I)** This cross-day backward shifting effect was found to be consistent across animals (plots show mean shift per animal, one-way boot-strap test of the mean, p < 0.006). These results suggest the BTSP-like mechanism may also play a role in cross-day, as well as within-session, spatial coding dynamics.

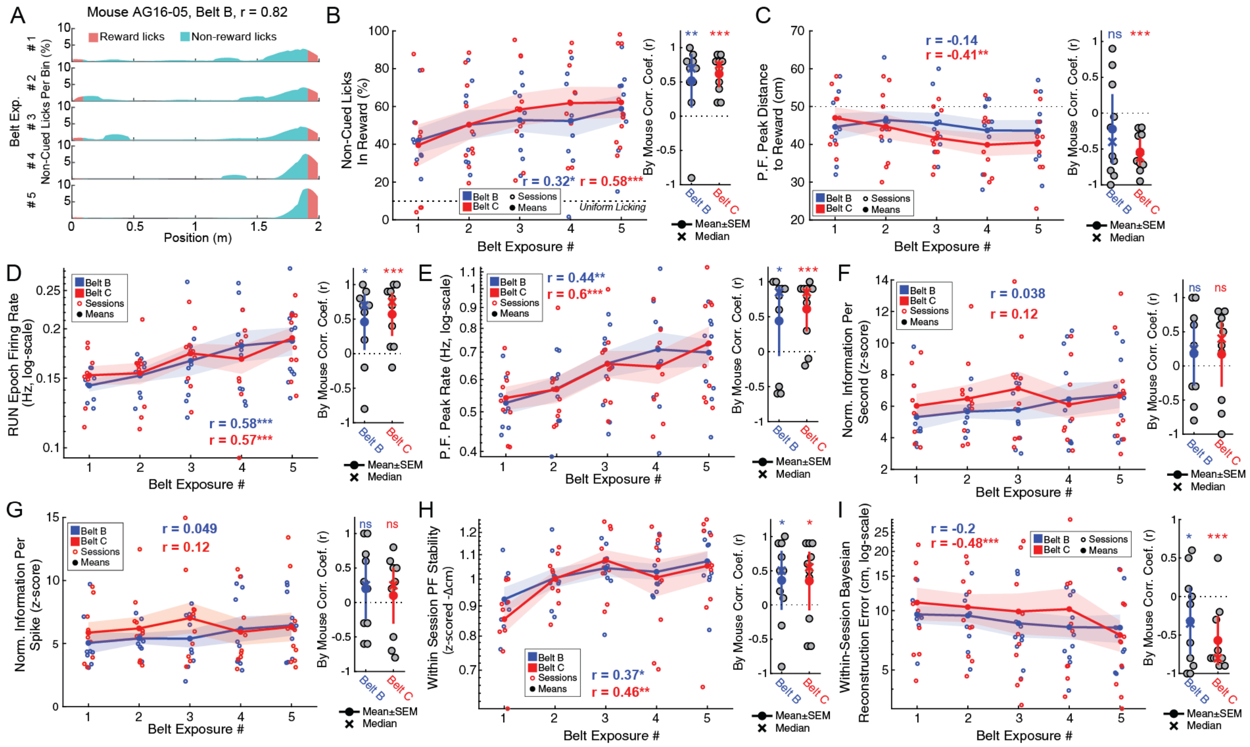


**Fig. S5: Evolution of behavior and hippocampal spatial coding over sequential exposures to the RUN belts.** The mice were exposed to each of the two RUN belts, B and C, for a total of 5 days each. The behavioral task required the animals to lick within a hidden rewarded zone to receive water reward. When the animals licked inside the reward zone (either due to learning or by chance) they received water which consistently elicited a 'cued', therefore putatively non-learning-related, licking bout. Therefore, in order to analyze the animal's learning-related behavior we excluded 'cued' licks occurring up to 1 second during or after water delivery ('non-cued licks'). **(A)** The percentage of non-cued licks by bin (100 bins total) are shown for each of the 5 days one mouse ran on Belt B, non-cued licks on the reward zone are colored in red. Note that the animal suppresses outside of reward licking as it gains experience over days on the belt. **(B,** left panel) The percentage of within-reward zone non-cued licks is shown as a function of RUN belt (blue and red for belts B and C, respectively) and belt exposure number (the number of sessions the animal had been run on the belt, solid circles/lines shown by belt means, shaded regions show ±SEM). The animals showed increasingly selective within-reward zone licking as they became familiarized with the belt. (B, right panel) the Pearson's correlation coefficient between belt exposure number and percentage of non-cued licks in the reward zone is shown per animal per belt, confirming this effect is consistent across animals (one sided boot-strap tests of the mean: belt B: $p < 0.0038$, belt C: $p \sim 0$). **(C)** The mean PF distance to reward is plotted per animal per belt as in (B, dashed horizontal line shows expected PF reward distance of uniform spatial tiling). While the reward zone was generally over-represented (sessions below the dashed line), consistent with previous reports [4,5] reward-related enrichment occurred more robustly on the second belt the animals were exposed to (belt C) as compared to the first (belt B, per animal per belt correlations for belt B: $p < 0.14$, belt: C $p \sim 0$). **(D)** The overall firing rates of place cells during online running bouts during the RUN epoch as well **(E)** their peak within PF firing rates increased significantly over repeated belt exposures. However, neither the mean shuffle-normalized spatial information per second **(F)**, or per spike **(G)** showed a clear correlation with repeated RUN belt exposures. Conversely, the within session stability (first to last half of laps spatial coding stability) increased significantly over repeated belt sessions (per animal per belt correlations for belt B: $p < 0.029$, belt C: $p < 0.032$). Therefore, while place cells do not become significantly more spatially informative over time, their coding does become more reliably stabilized to a location. **(I)** As a consequence, Bayesian decoding of position generally becomes more reliable with experience (plots show median absolute Bayesian reconstruction error across sessions (per animal per belt correlations for belt B: $p < 0.04$, belt C: $p < 2.6x10^{-4}$)).

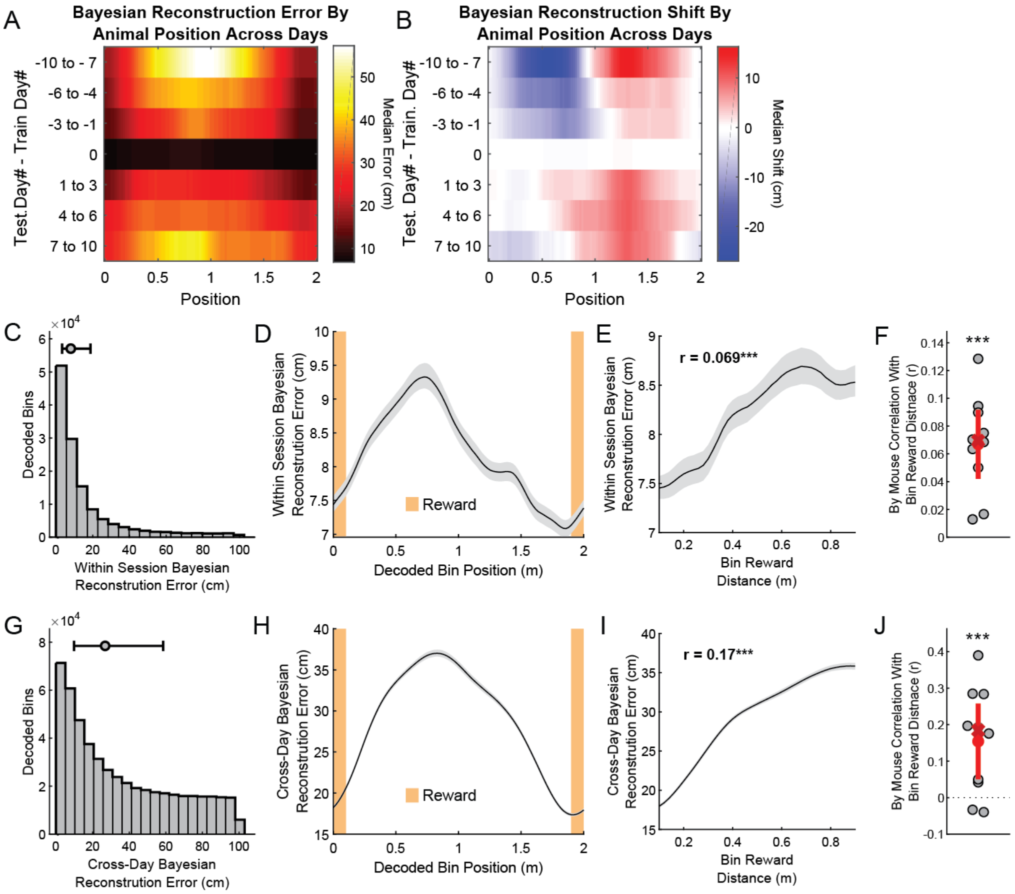


**Fig. S6: Within and across day Bayesian population decoding varies systematically with distance to reward.** **(A)** A heat map showing the median decoding error as a function of the animal's physical position on the RUN belt within (middle row) or across days (other rows, same-belt cross-day comparisons only, medians taken within a ±20 cm sliding window across the belt (x-axis)). Note that decoding error tends to be largest when the animal is far from the reward. **(B)** Median decoder 'shift' (the signed difference between the decoded and observed positions) is plotted as a function of the animals' physical location on the RUN belt as in (A). Note that, consistent with the observed backward shift of place fields over days (Fig S4, H, I) when past day's place fields are used as a template to decode the animal's position on future day's (last three-rows, positive values of train – test day) the decoder tends to predict positions ahead of the animal's physical location (red heat-map values). **(C)** The observed per bin distribution in cross-validated within-session error is shown (n = 133,276 third of a second RUN bins, horizontal bars show the median and $25^{th}$ to $75^{th}$ percentiles). **(D)** Within-session reconstruction errors are plotted as a function of observed (physical) position along the RUN belt (plots show medians ±boot-strapped 95% confidence interval across bins in a circularly sliding ±20 cm window). **(E)** Within-session reconstruction errors are shown as a function of each bin's observed distance to reward (plots show cross-bin in medians ±boot-strapped 95% confidence interval in a sliding 20 cm window, inset shows the correlation coefficient between bins reward distance and reconstruction error, p ~ 0). **(F)** The within-mouse correlation between bin reward distance and reconstruction error is shown (p-value from one-sided boot-strap test of the mean correlation coefficient, p ~ 0, n = 10 mice). **(G)** The distribution of same RUN belt, cross-day Bayesian reconstruction errors is shown as in (C, n = 508,779 cross-day decoded RUN bins). **(H)**

Cross-day reconstruction errors are plotted as a function of observed position (plotted as in D). **(I)** Cross-day reconstruction errors are plotted as a function of observed distance to reward as in (E, $p \sim 0$). **(J)** The within-mouse correlation between distance to reward and reconstruction error is plotted as in (F, $p < 6x10^{-5}$) revealing that the overall higher population decoding accuracy observed near the reward locations is consistent across animals.

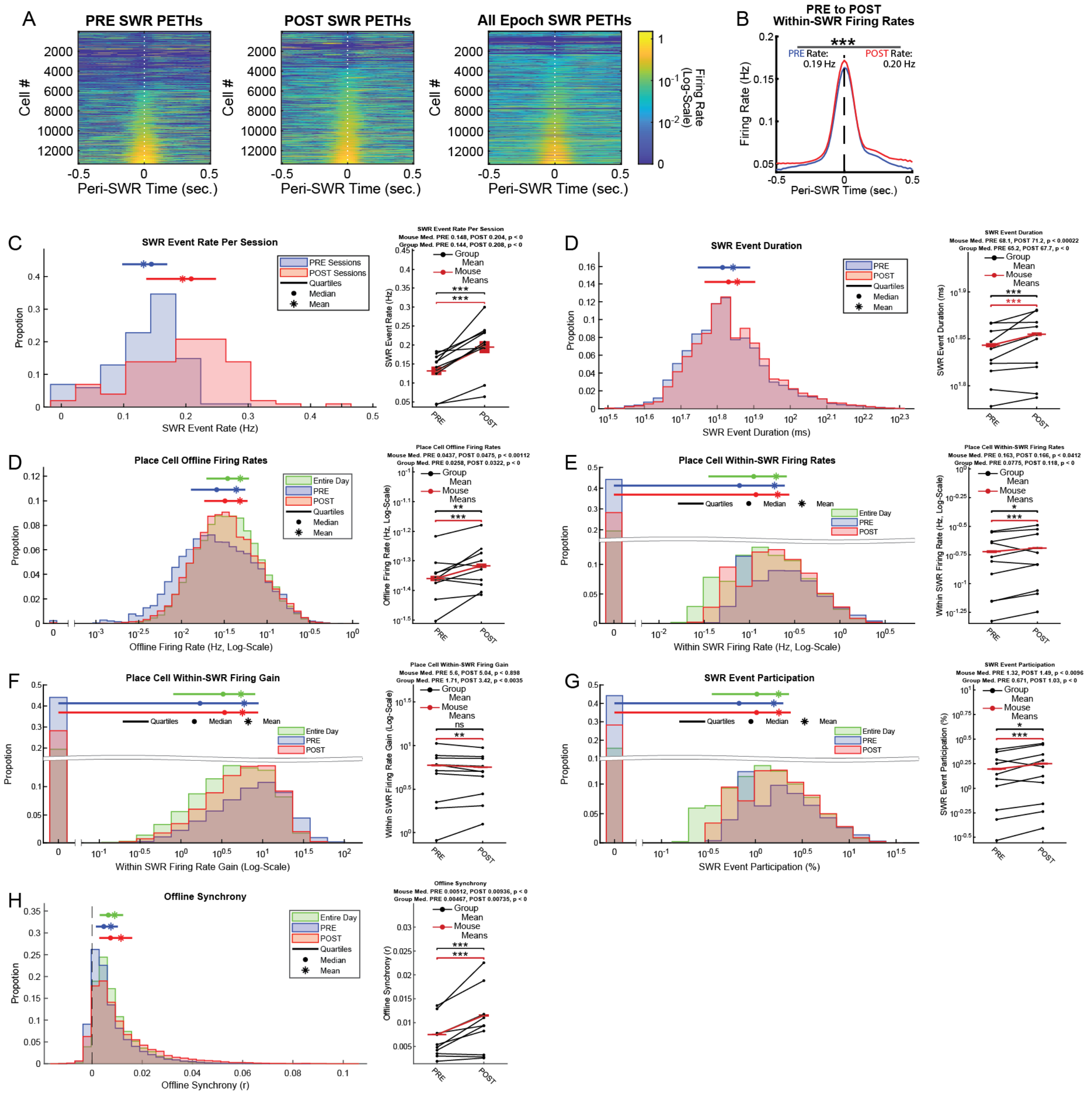


**Figure S7: Offline activity increases from the PRE to the POST epoch.** **(A)** Peri-event time histograms (PETHs) were made around SWR peaks for each place cell on each imaging day (rows), either for the PRE epoch (left panel), the POST (right panel), or for the entire day's offline activity combined. Cells are arranged from lowest to highest within-SWR firing rates in each epoch. Note that most place cells show a sharp positive modulation by SWR's during the offline state (10,194/13,341 place cells (76.4%) fired in at least one SWR event throughout the day, and 9,707 place cells (72.7%) showed greater within SWR than overall immobility firing rates). **(B)** Mean peri-SWR PETH's for the POST and the PRE epoch showing a small but significant increase in the within-SWR firing rates from PRE to POST (PRE mean: 0.189 Hz, median: 0.077 Hz, POST mean: 0.204 Hz, median: 0.118 Hz, Signed-Rank p ~ 0, n = 13,341 place cells). **(C,** left panel**)** The distribution offline SWR event rate per session (total SWR events/total offline seconds) is plotted for PRE (blue) and POST (red) sessions. (C, right panel)

The per session SWR event rates are plotted for the group data (red lines, vertical red bars show the group mean ± SEM, n = 101 PRE and POST sessions each, Signed-Rank p ~ 0) and the mean for individual animals (n = 10 mice, paired one-way boot-strap test of the mean, p ~ 0). **(D)** Likewise, the average duration of SWR events increased from the PRE to the POST epoch (group PRE mean: 68.8 ms, median: 64.0 ms, POST mean: 70.7 ms, median: 67.2 ms, Ranked-Sum test p ~ 0, n = 31,676 SWR events). **(D)** Overall place cell offline (immobility) firing rates decreased significantly from the PRE to POST epochs (group PRE mean: 0.043 Hz, median: 0.025 Hz, POST mean: 0.048 Hz, median: 0.032 Hz, Signed-Rank test p ~ 0, n = 13,341 place cells). **(E)** Within SWR firing rates also increased significantly from the PRE to the POST epoch (group PRE mean: 0.189 Hz, median: 0.077 Hz, POST mean: 0.205 Hz, median: 0.118 Hz, Signed-Rank test p ~ 0, n = 13,341 place cells). **(F)** The median SWR-firing rate gain (within SWR firing rate divided by overall offline firing rate) increased significantly across cells grouped across animals (group PRE mean: 5.70, median: 1.67, POST mean: 5.63, median: 3.41, Signed-Rank test p < 0.0034, n = 13,341 place cells), however this effect varied across mice (right panel, black lines). **(G)** Furthermore, place cells tended to participate (i.e. have at least one calcium estimated spike occurring within) a greater percentage of POST as compared to PRE SWR events (group PRE mean: 1.57%, median: 0.67%, POST mean: 1.78%, median: 1.03%, Signed-Rank test p ~ 0, n = 13,341 place cells). **(H)** Together these PRE to POST activity changes led to a robust increase in per cell population offline synchrony calculated as the mean of the Pearson's correlation coefficient of each place cell's offline firing rate vector (smoothed with 150 ms Gaussian kernel) with that of every other place cell (group PRE mean: 0.007, median: 0.005, POST mean: 0.012, median: 0.007, Signed-Rank test p ~ 0, n = 13,341 place cells). Similar, though slightly smaller magnitude, effects were observed when all (including non-place cell) ROI's were considered (data not shown).

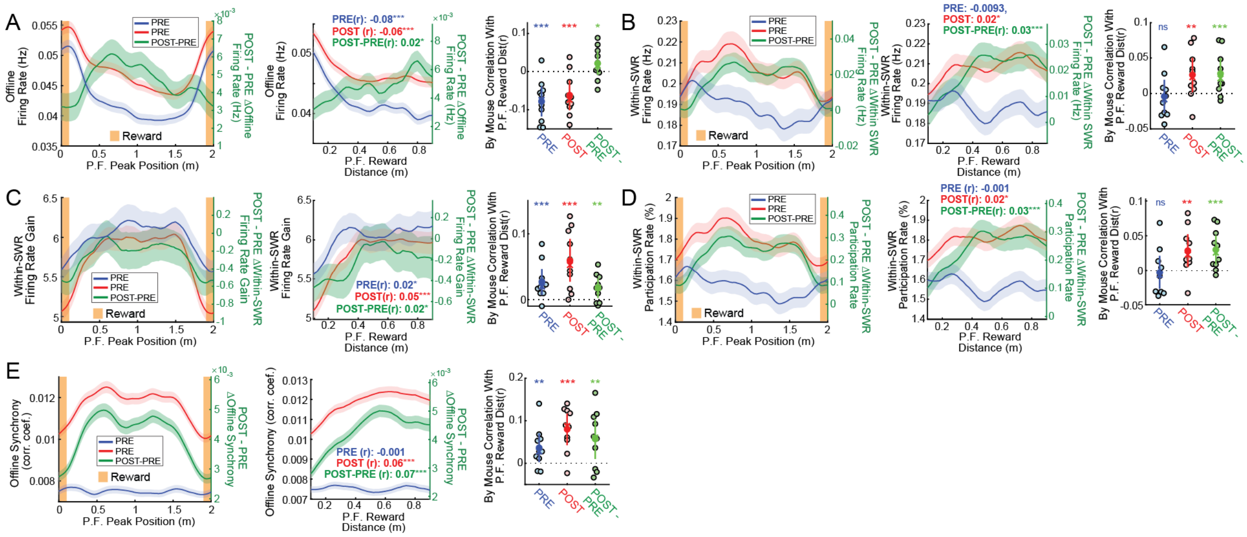


**Fig. S8. PRE to POST changes in calcium offline activity and synchrony correlate with place field distance from the reward during the RUN.** **(A,** left panel**)** Place cell offline (immobility) firing rates are plotted for either the PRE (blue) or POST (red) within a circularly sliding ±20 cm window of their PF peak position during the intervening RUN (left-ordinate, mean±SEM). Overlaid are the by-position POST-PRE differences (green, right ordinate). (A, middle-panel) PRE, POST (left ordinate) and POST – PRE (right-ordinate) offline firing rates are plotted as a function of PF peak distance to reward in a ±20 cm window. Inset numerical values show the Pearsons correlation coefficients between the group offline firing rates and RUN PF peak distance to reward revealing that while offline firing rates tend to be larger for near-reward coding cells both during PRE and POST (significant negative correlations, Fisher's z-test, *p<0.05, **p<0.005, ***p<0.0005, n = 13,341 place cells), increases in PRE to POST offline firing rate are largest for cells coding far from the reward (green inset, small but significant positive correlation). (A, right panel) These patterns were confirmed by a calculating the per mouse Pearson's correlation coefficients between PRE, POST or POST – PRE offline firing rates to RUN PF peak reward distance (y-axis, p-values from one-sided boot-strap test of the mean, solid circles and vertical lines show mean ±SEM across mice, crosses show across mouse median correlation coefficient values). **(B)** Within-SWR firing rates are plotted as in panel A, revealing that while no correlation with reward distance were observed for within-SWR firing rates during the PRE epoch, both the POST epoch and POST – PRE difference in within-SWR firing rates are correlated with RUN PF distance to reward within and across animals. Consistent with this finding the POST epoch and POST-PRE differences in both **(C)** the within SWR-firing rate gain and **(D)** the percentage of SWR's each cell participated in, were positively correlated with RUN PF distance to reward across cells within and across animals. **(E)** Together these effects led to a larger increase in by cell population synchrony for cells father from the reward than cells close to the reward.

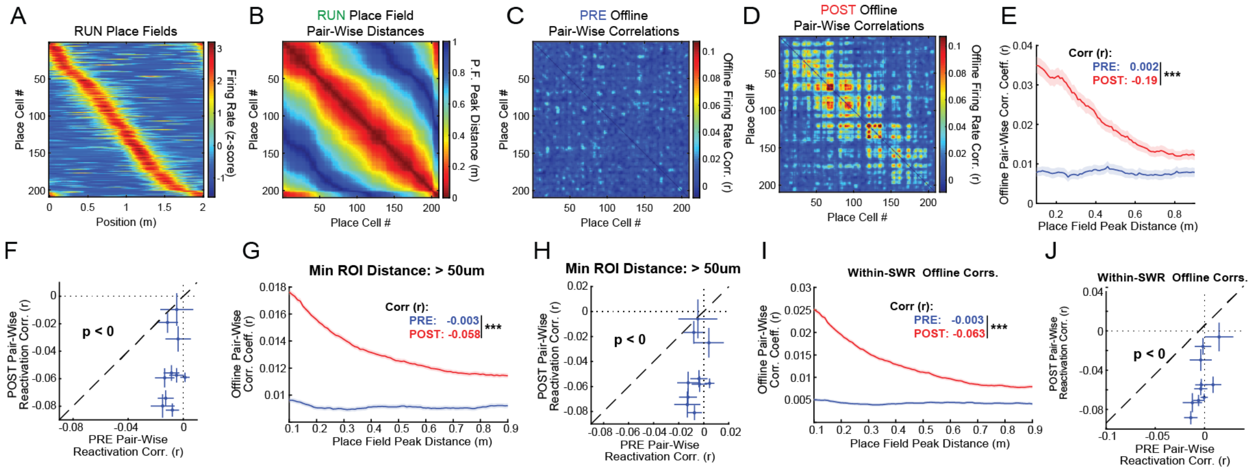


**Fig. S9: Pair-wise place field distance relationships are reactivated during the POST offline epoch.** **(A)** The z-scored firing rate by position vectors (sorted by PF peak) are shown for a representative RUN session. **(B)** These were used to compute a pair-wise circular PF-peak distance matrix. The pair-wise offline firing-rate correlations (Pearson's correlations between the immobility firing rate vectors convolved with 150 ms Gaussian kernel) are shown after being smoothed with a 2d 2.5 cell Gaussian kernel for illustration clarity for either the **(C)** PRE or **(D)** POST offline epochs (same place cell correlations were removed from the analysis). Note that cells with nearby place fields during the intervening RUN show elevated correlations during the POST as compared to PRE epochs. **(E)** The reactivation relationship for this example day is shown by plotting the mean pair-wise offline correlation coefficents (y-axis) in a sliding 10 cm window of PF peak pairwise distances (x-axis, shaded regions show the boot-strapped 95% confidence interval). Insets show the correlation coefficient between offline pair-wise correlations and pair-wise PF peak distances showing that cells with nearby place fields peaks tend to be more correlated during the POST than during the PRE (inset p-value, Fisher Z-test, $*p<0.05$, $**p<0.005$, $***p<0.0005$). **(F)** The pair-wise reactivation effect observed for the group data (Fig. 2E) was replicated on a per-mouse basis. Plots show the per mouse correlation coefficients of RUN PF peak distance to offline pair-wise correlations either during the PRE (x-axis) or POST (y-axis) epochs (bars show 95% confidence interval, one-sided boot-strap test of the mean correlation coefficient, $p \sim 0$, $n = 10$ mice). **(G)** In order to control for possibly spurious local correlations in the field of view (FOV), the pair-wise correlation analysis was repeated after eliminating pairs of cells with ROI pair-wise center of mass distances of less than 50 μm. This analysis as well as its by-mouse variant **(H)** confirmed that the POST increase in offline correlation of cells with nearby PF's is a global, rather than strictly local effect. **(I, J)** Finally, carrying out the pair-wise reactivation analysis only including within-SWR offline epochs (1.17% of total offline time) preserved the pair-wise reactivation effect, suggesting SWR-related activity is a major driver of the pair-wise reactivation signal.

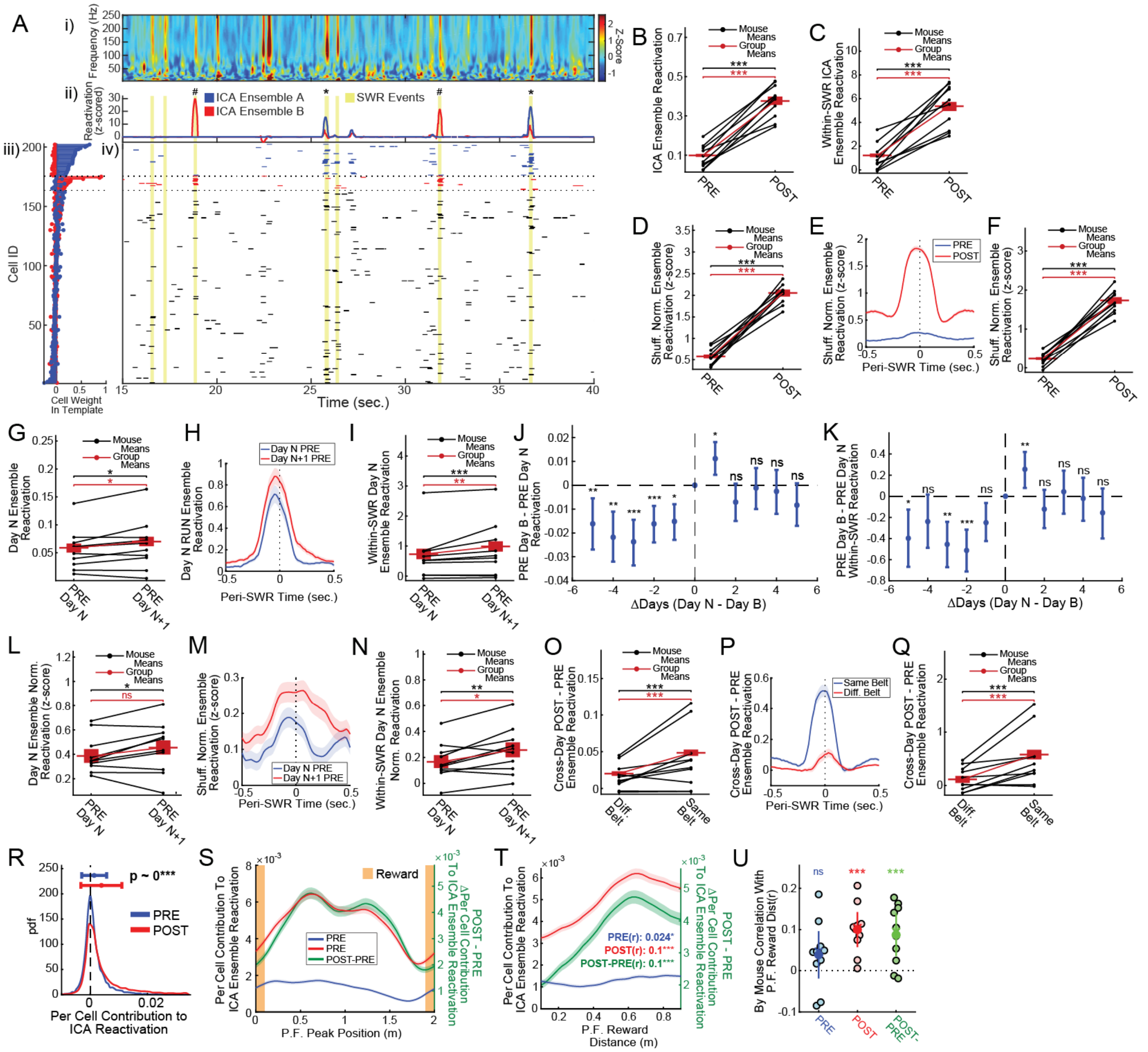


**Fig. S10: Offline activity shows belt-specific long-lasting reactivation of RUN-related ICA ensembles.** **(A)** A representative example from one POST session showing the wavelet spectrogram (panel i) and reactivation time-courses for two RUN-related ICA ensembles (panel ii, red and blue lines, LFP-detected SWR events are highlighted in yellow). Panel iii shows each place cell's ICA component weight for each of the two ensembles with dashed lines separating those cells strongly recruited to ensemble A (top), ensemble B (middle), or neither (bottom, for the purposes of this illustration strongly contributing cells were those with ≥1S.D. above the mean for each template; no cells were strongly modulated in both templates in this example). Panel iv shows the offline firing rate raster (convolved with a 150 ms Gaussian kernel). Note that ensemble B is strongly recruited to the third and sixth LFP-detected SWR-events in this period (marked with a '#' symbol, panel ii, top), while ensemble A is strongly recruited to the fourth and seventh SWR-events detected in this epoch (marked with a '*' symbol), demonstrating the robust SWR-modulation and ensemble specific reactivation of RUN-related ICA ensembles during offline epochs. **(B)** The PRE to POST increases in RUN ensemble offline reactivation were confirmed across mice (black lines: per animal means, p-value from one-sided boot-strap test of the mean difference, n = 10 mice; red lines: group data, vertical line shows group mean, horizontal red bars show ±SEM, p-value from Signed-Rank test of per RUN ensemble POST – PRE reactivation differences, n = 3,843 RUN ensembles). **(C)** A similar per animal analysis was carried out for the within-SWR reactivation values. **(D)** In order to account for none-specific changes in excitability and synchrony from the PRE to the POST epoch 1,000 null reactivation scores were computed for each ensemble by randomly permuting the template cell id's – each cell's observed reactivation value was normalized (z-scored by) its null distribution such that its expected null value was 0. The robust increase in offline POST as compared to offline PRE shuffle-normalized RUN-ensemble reactivation suggests that the reactivation effect was not attributable to non-specific excitability or synchrony changes. **(E)** The peri-SWR PETH's of shuffle normalized RUN-ensemble reactivation (±SEM) were constructed, confirming that RUN-specific content increases robustly around the time of SWR's. **(F)** This within SWR-shuffle normalized PRE to POST increase was confirmed on a per-mouse basis. **(G)** In order to examine the duration of RUN-ensemble reactivation day *N* RUN-ensembles were used to measure either to Day *N* PRE (~20 minutes earlier) or Day *N+1* PRE (~24 hours after) the RUN. Note that for all multi-day comparisons ensembles were recomputed using only the subset of cells which were place cells on the template day and which were registered on both days of the comparison – consequently the templates applied were identical on both days of the comparison. Notably, offline reactivations of RUN day *N* ensembles were larger on PRE Day *N + 1* compared to PRE day *N*, an effect observed both at the group level and across mice. **(H)** This effect was found to be enhanced around the time of PRE SWR's and was **(I)** significant at the group and by-mouse levels within-SWR epochs. Extending the cross-day analysis to all pairs of Day *N*, versus day *B* PRE epochs revealed that the reactivation effect is specific to the Day *N+1* time point both for overall offline activity **(J)** and for within-SWR activity **(K)**, suggesting that though occurring over much longer time-courses (~1 day) than the previously reported minutes-long decay [6] though see [7,8]) reactivation nonetheless decays over time. While the combinatorial explosion associated with the all-day-to-all-day comparison shown in panels *J & K* make calculating the shuffled reactivation scores computational unfeasible, we none-the-less applied this analysis to the PRE day *N* to PRE day *N+1* comparisons. **(L)** This revealed that the entire offline epoch shuffle-normalized day *N* vs. *N+1* effect is not quite robust enough to be distinguished from chance at the group level (PRE day *N* mean: 0.38, median: 0.20, PRE day *N+1* mean: 0.39, median: 0.27, Signed-Rank $p < 0.052$, n = 1,750 ensembles). **(M)** However, the shuffle-normalized PRE day *N* vs. *N+1* effect was enriched around the time of SWR's, and **(N)** was significant at the groups and animal levels when restricting the analysis to within-SWR epochs. **(O)** In order to assess the specificity of ICA ensemble reactivations to given belts, day *N* RUN ensembles were used to assess POST – PRE changes in reactivation across days either when the animal was run on the same RUN belt (belt B to belt B or belt C to belt C, comparisons) or the other RUN belt (belt B to belt C or belt C to belt B). It was found that both at the group and animal levels PRE to POST increases in reactivation were larger when the two days being compared were run on the same RUN belt. **(P)** This effect was largest around the time of SWR's and **(Q)** was significant at the group and animal level when considering only PRE to POST changes in within-SWR activity. Per cell contribution (PCC) to ensemble reactivation was estimated as the mean across ensembles of the observed offline reactivation (as in panel B) minus the reactivation observed after repeating the analysis with that place cell left out. **(R)** As expected PCC's were higher in the PRE as compared to POST epochs (plots show the distribution of PRE and POST PCC values, PRE mean: $1.28 \times 10^{-3}$, median: $4.8 \times 10^{-4}$, POST mean: $4.7x10^{-3}$, median: $1.5x10^{-3}$, Signed-Rank p ~ 0, n = 13,341 place cells). **(S)** PCC values (left axis) and PRE to POST changes in PCC values (right axis) are plotted as a function of place cell's PF peak position during the RUN (±20 cm sliding window, mean ±SEM, inset values shows the Pearson's correlation coefficient between PCC value and PF distance to reward) revealing that place cells coding further from the reward preferentially contribute both POST epoch PCC scores and the PRE to POST changes in PCC scores as compared to PRE PCC scores, **(U)** an effect which was confirmed by a within-animal correlation analysis.

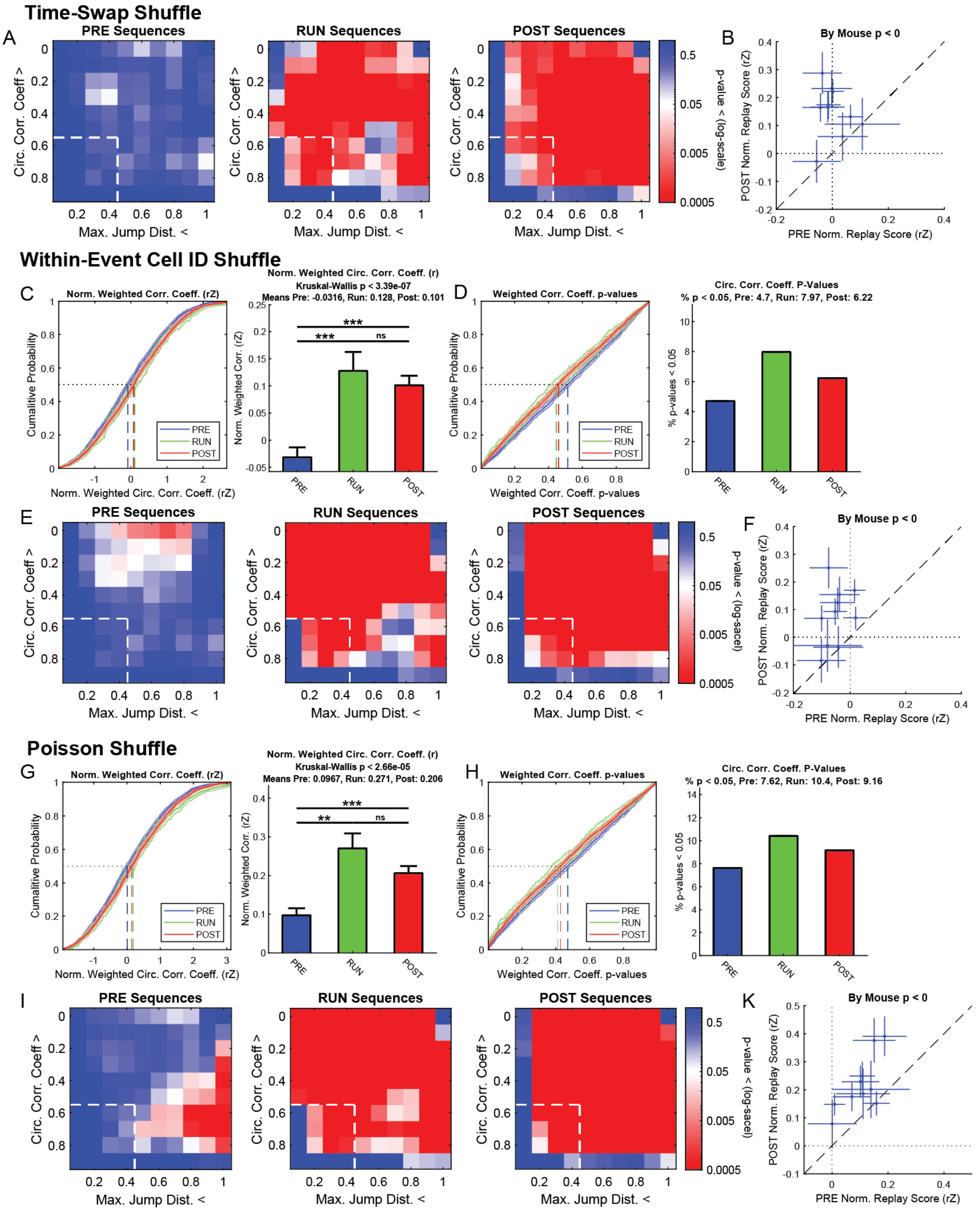


**Figure S11: Calcium estimated sequence replay is statistically robust.** Within-PSE sequence content was estimated from the estimated within PSE-event Bayesian posterior probabilities using a version of the weighted-correlation method [9] adapted for circular trajectories. This method was validated by visual examination as well as by comparison to the older 'line-casting'

(or 'Radon' transformation) technique which led to similar results (data not shown). The principal shuffling method employed in the study is the temporal bin shuffle in which the order of the bins within each event is permuted (resampled without replacement) 2,000 times and the identical sequence replay analysis is conducted of each of these shuffled events. Because differences in event activity and duration can lead large differences in each event's expected null distribution [8] each event's observed sequence score (r) was normalized (z-scored by) their respective shuffled distribution (rZ score). Time-swap shuffle rZ scores were to found to increase significantly from the PRE to POST (Fig 3C, rZ PRE mean: 0.003, median: -0.04, POST mean: 0.17, median: 0.13, Ranked-Sum test p ~ 0, n = 2,977 PRE and 3,437 POST PSE's). **(A)** Following [10] the observed sequence content was further verified by jointly comparing the observed sequence score and maximum jump distance (the maximum absolute circular distance between the reconstructed Posterior probability peaks on the successive bins of the event) to those observed in the shuffle distribution. The panels show the empirical p-value (the proportion of shuffled datasets in which at least the same number or more events met the inclusion criteria as the number of events meeting criteria observed in the experimental dataset) for the sequence score and maximum jump distance thresholds indicated on the y and x axes respectively (heat-map show p-value on a log-scale, red values indicate that the experimental data sets show significantly more events passing criteria than would be expected based on the shuffled datasets, dashed lines show the 'virtual trajectory' criteria boundary suggested by Silva et. al. 2015). **(B)** PRE to POST increases in shuffled normalized sequence scores (rZ) were consistent across mice (plot shows mean ±SEM, one-sided boot-strap test of mean difference p ~ 0). **(C)** Given that each type of shuffle makes subtly different assumptions [8,11] we also tested out dataset using an alternative shuffling procedure in which the cell ID's of cells active (firing at least one estimated spike) in each event were shuffled within-event, preserving each event's mean firing rate structure as well as the number of events each cell participated in, though not each cell's firing rate or firing-structure in each event. **(C,** right panel**)** The cumulative distribution of shuffle normalized sequence scores during offline PSE's occurring during the PRE, RUN or POST epochs is shown (vertical lines show distribution medians) as well as the mean (±SEM) for each epoch (C, right panel, Kruskal-Wallis $p < 3.4 \times 10^{-7}$). **(D)** The cumulative distribution of per PSE p-values (left panel) and percentage of significant event is shown (at $p < 0.05$, PRE: 4.7%, RUN: 8.0%, POST: 6.2%). **(E)** The joint circular weighted correlation coefficient and maximum jump distance analysis was performed as in panel A. **(F)** The per animal increases in PRE to POST sequence content were confirmed as in panel B. Finally, generating synthetic datasets from Poisson distributions matching each cell's per epoch within PSE firing rate has been proposed as a surprisingly stringent null hypothesis model to test sequence content [10]. Though there are some concerns whether this is indeed a valid null hypothesis set (see [8]) out of an abundance of caution we nevertheless tested our data against this null. Panels **(G, H, I** and **K)** are plotted as in C, D, E and F, respectively. These results confirm the accessibility of canonical sequential virtual hippocampal trajectories to fast calcium imaging methods.

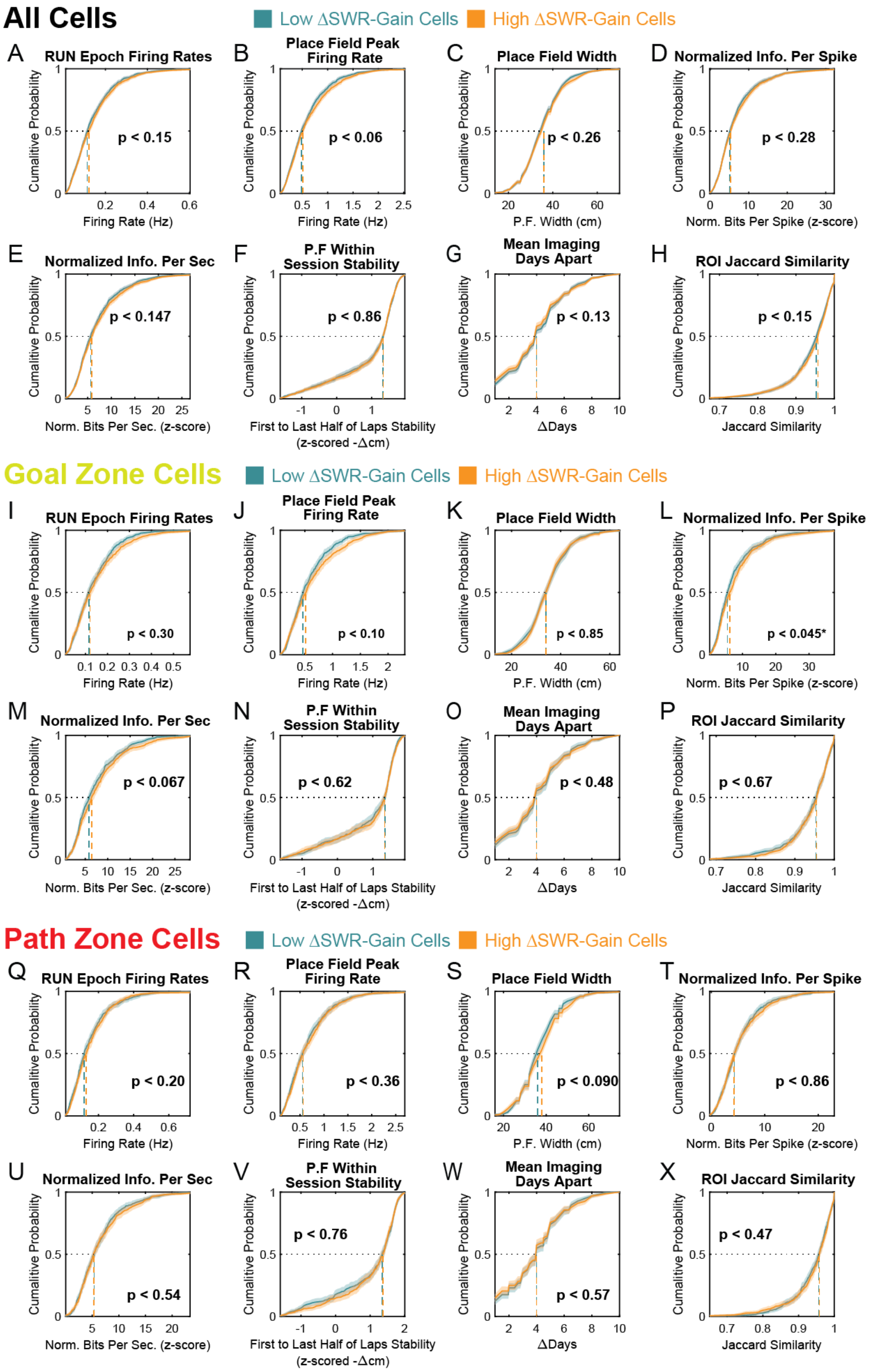


**Fig. S12: Spatial coding properties of Low and High ΔSWR-Gain cells do not differ on the novel place field day.** For the per-cell stability analysis (Fig. 4, A-G) for a given day place cells which showed their first place field on that day (novel place field day) for a given RUN belt (B or C) were grouped by their PRE to POST changes in SWR-gain. It was found that High ΔSWR-Gain cell's novel place fields were more stable as assessed by the mean of their subsequent day's RUNs on the same belt for which the same cell also showed significant spatial coding. This insured that each cell contributed to the analysis at most twice (once for each belt). However, the observed future stability of the High ΔSWR-Gain cell group could be controlled by spatial coding differences during the RUN on the novel place field day. Therefore, we compared the spatial coding properties of cells Low and High ΔSWR-Gain cell groups for the novel place field day. **(A)** Overall firing rates during novel place field day running bouts did not differ between the Low and High ΔSWR-Gain cell group (graphs show cumulative distributions ±boot-strapped 95% confidence interval, dashed vertical lines indicate the group medians, Low Group mean: 0.14 Hz, median: 0.11 Hz, High Group Mean: 0.15 Hz, median: 0.12 Hz, Ranked-Sum test $p < 0.15$, $n = 1{,}212$ novel place cells per group). **(B)** Their place field peak rates did not differ significantly (Low Group Mean: 0.6 Hz, median: 0.48 Hz, High Group Mean: 0.64 Hz, median: 0.51 Hz, $p < 0.061$). **(C)** The width of their place fields was likewise similar between groups (Low Group Mean: 36 cm, median: 36 cm, High Group Mean: 37 cm, median: 36 cm, $p < 0.26$). **(D)** They did not differ in shuffle-normalized spatial information per spike (Low Group Mean: 6.7, median: 5.1, High Group Mean: 6.9, median: 5.2, $p < 0.29$), nor **(E)** normalized information per second (Low Group Mean: 6.9, median: 5.6, High Group Mean: 7.3, median: 5.8, $p < 0.15$). **(F)** The groups also did not differ in their novel place field day within session stability (first half of laps to second half of laps, Low Group Mean: 0.95, median: 1.3, High Group Mean: 0.96, median: 1.3, $p < 0.86$). **(G)** Since in the per cell analysis (Fig 4A-G) cells were compared to the mean of their future stability, it could be the case that, on average the comparison days (i.e. future days that cell had a place field on that belt) occurred closer or further in time between the groups, however, a comparison the difference between the novel place fields days and the mean comparison days showed the two groups did not significantly differ (Low Group Mean: 4.2 ΔDays, median: 4 ΔDays, High Group Mean: 4.1 ΔDays, median: 4 ΔDays, $p < 0.13$). **(H)** Finally the cross-day registration quality (the mean of the pair-wise Jaccard similarity scores of the novel place field days and the future place field days) was compared between groups, showing no significant differences (Low Group Mean: 0.94, median: 0.95, High Group Mean: 0.94, median: 0.96, $p < 0.15$). **(I-P)** The analysis outlined in panels A-H was applied to the goal-zone cell group ($n = 513$ novel place fields per group). **(L)** While a small difference was observed in goal-zone cell's shuffle-normalized information per spike (Low Group Mean: 7.4, median: 5.5, High Group Mean: 8.1, median: 6.2, $p < 0.045$) this relationship does not logically help to explain the observed lack difference in the future stability of Low and High ΔSWR-Gain cells (Fig. 4G) observed in the goal-zone group. **(Q-X)** No significant difference were observed between the path-zone Low and High ΔSWR-Gain groups (plotted as in A-H).

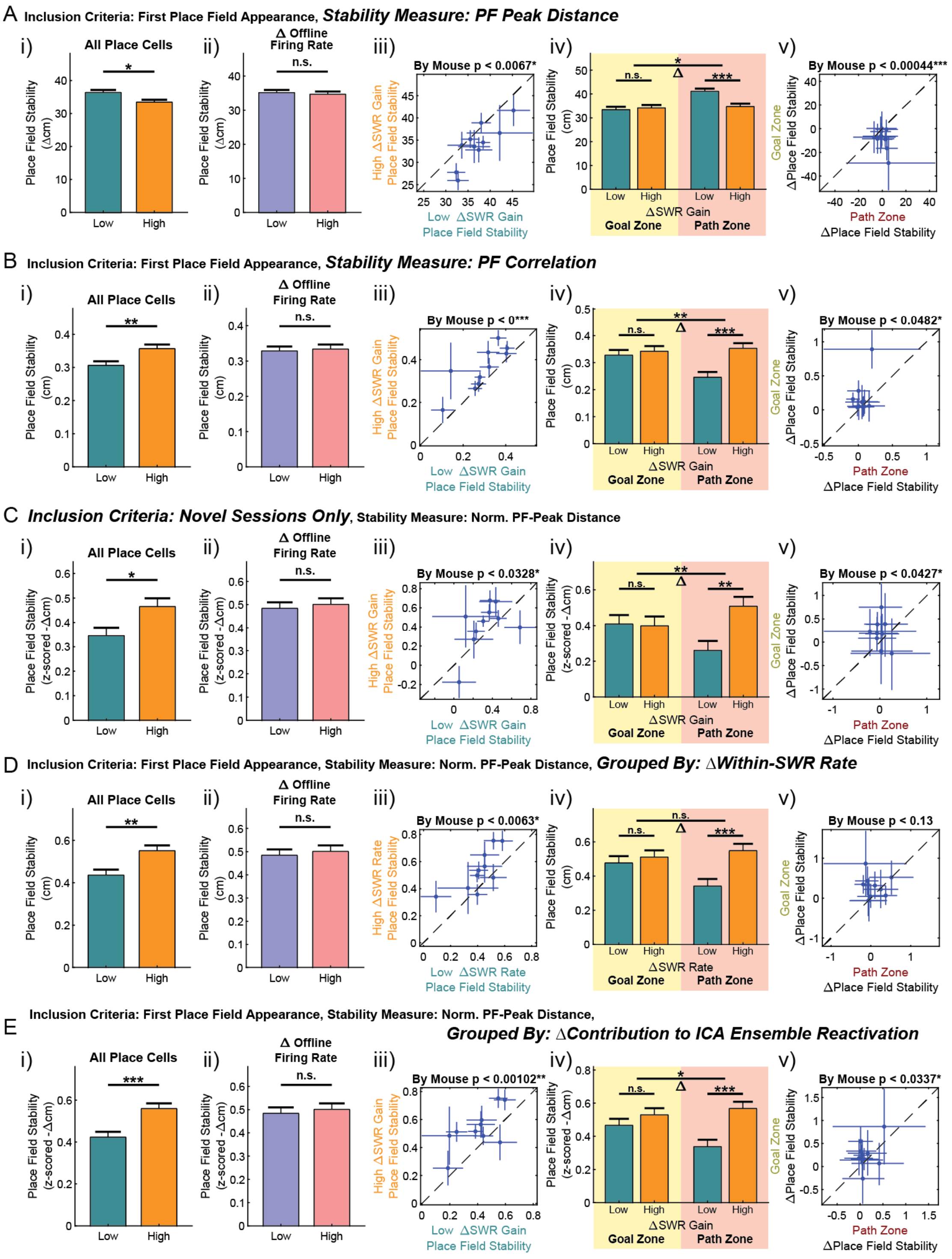


**Fig. S13: PRE to POST change in SWR-specific recruitment reliably predict increase path zone cell spatial stability.** In order to test whether the observed relationship between PRE to POST changes in SWR-gain and future place field stability, and its difference between the Goal and Path zone cell groups (Fig. 4, A-G), were robust across conditions, parallel analyses were carried out across a variety of methods for measuring cross-day stability, different cell inclusion criteria and measurements of SWR-recruitment. **(A)** Low and High ΔSWR-gain groups were identified as in Fig 4A, however, the non-normalized PF peak distance was used as a measure of stability. (A, panel i) Under this stability measurement novel place field day High ΔSWR-gain predicted novel place field stability (note that in this measurement lower values correspond to more stable cells, p-value from Ranked-sum test, n = 1,212 novel place cells per group, *$p<0.05$, **$p<0.005$, ***$p<0.0005$ for all panels). (A, panel ii) However, no effect was observed when cells were instead grouped by their overall offline-firing rate changes from PRE to POST. (A, panel iii) The increased future stability of High ΔSWR-gain cells was consistent across animals (plot shows the within animal mean ±SEM for Low and High ΔSWR-gain groups in the x and y axes respectively, p-value shows from the one-sided boot strap test of the mean difference between groups across animas, n = 10 mice). (A, panel iv) Cells were separated into path and goal cells group as in Fig 4G. High ΔSWR-gain cells showed increased future stability specifically in the path zone group (Ranked-sum test, n = 513 and 509 novel place cells per Low/High group in the path zone and goal zone cell groups respectively). The significance of the interaction between the Path/Goal zone cell group, Low/High ΔSWR-gain cell group and future place field stability was computed via a permutation test in which the observed difference in Path cell [High – Low future stability] versus Goal cell [High – Low future stability] was compared to the same value computed in 25,000 permuted data sets in which High vs. Low ΔSWR-gain cell labels were randomly permuted (significance bar above the 'Δ' symbol)). (A, panel v) The interaction between Low vs. High ΔSWR-gain group and Path vs. Goal cell groups was found to be significant across mice (plots show the mean ±boot-strapped 95% confidence interval of the High – Low ΔSWR-gain group gain difference in future stability for the Goal (x-axis) and Path (y-axis) groups, p-value from one sided boot-strap test of the per animal mean Goal[High – Low] – Path[High – Low] difference across animals, n = 10 mice). **(B)** Consistent results were obtained when using pair-wise Pearson's correlations between RUN firing-rate by position vectors as a measure of stability instead of normalized PF distances (plotted as in A). **(C)** Next, we repeated the analysis performed in Fig 4A-G but restricting our analyses to only those novel place fields occurring during a novel RUN session (i.e. the first RUN session that the animal had run on that RUN belt, n=663 novel place fields per Low and High group). **(D)** We repeated the analysis shown in Fig 4A-G but we sorted the cells into equal groups based on their ΔSWR-rate (rather ΔSWR-gain). Though specific tests of Low/High ΔSWR-rate and goal vs path zone interactions were not significant under this condition (panels iv and v), this analysis further verified that PRE to POST changes in within-SWR recruitment predicts future place field stability (panels i and iii) and this this increase in stability is only significant for path zone cells (panel iv). **(F)** Finally, cells were split by cell's normalized PRE to POST change in contribution to ensemble (ICA) replay (as in Fig 4K), confirming that contribution to POST-specific RUN-ensemble reactivation predicts place cell's future day stability, specifically for Path zone cells.

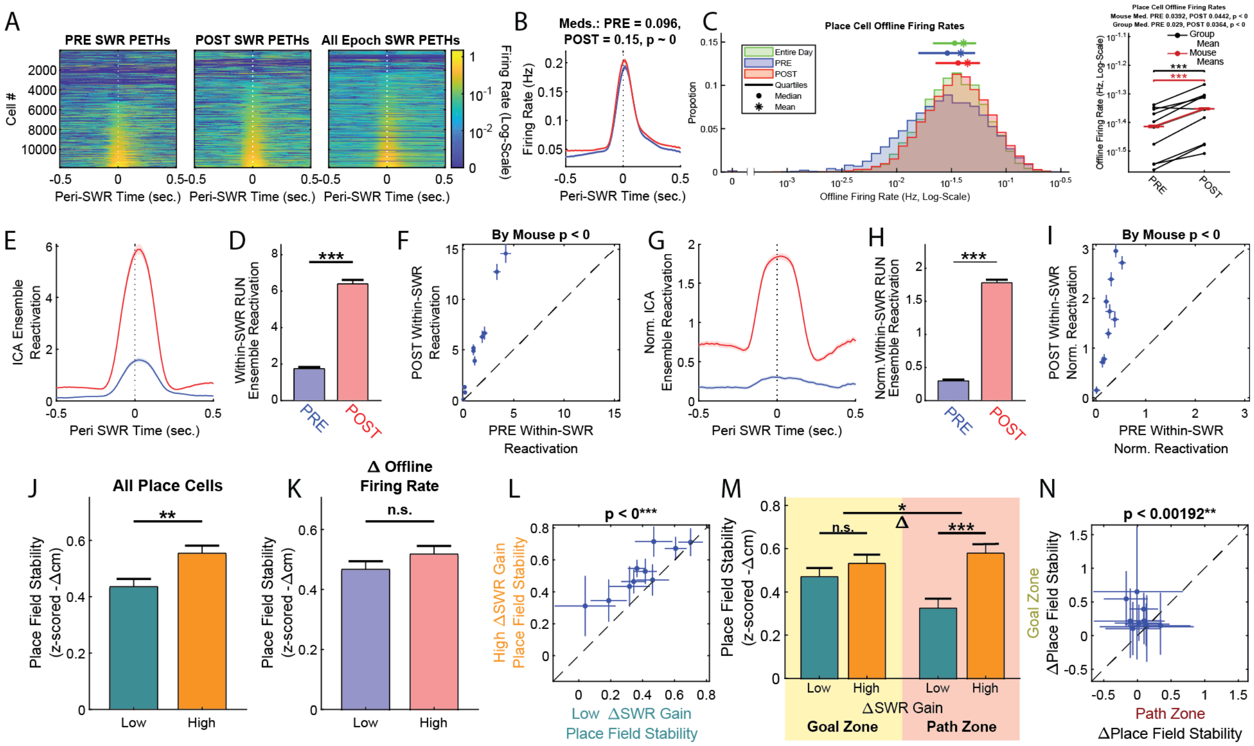


**Fig. S14: Transient-based analysis of offline activity, reactivation and SWR-related place field stabilization.** As an additional control to verify that our results could not be a consequence of the spike deconvolution approach used to estimate activity use elsewhere throughout this study, we additionally employed a transient-based analysis (see Methods). **(A)** The transient-based SWR-triggered PETH's are shown as in fig. S7A, demonstrating that offline transient rates are enriched within SWR-events (9,113/11,837 place cells (76.9%) displayed a transient in at least one SWR throughout the day, and 76.7% of place cells showed elevated within-SWR rates as compared to total immobility firing rates). **(B)** POST epoch within-SWR transients rates increased relative to PRE rates (PRE within SWR-transient rate mean: 0.23, median: 0.096, POST mean: 0.26 Hz, median: 0.14 Hz, Signed-Rank test $p \sim 0$, $n = 11{,}837$ place cells). **(C)** For comparison overall offline firing rates are plotted as in fig. S7D (PRE mean: 0.039 Hz, median: 0.029 Hz, POST mean: 0.044 Hz, median: 0.036 Hz, Signed-Rank $p \sim 0$). **(D)** A peri-SWR transient-based ICA RUN-ensemble analysis was carried out as in Fig 2G. **(E)** Both overall (data not shown) and within-SWR RUN ensemble reactivation increased from PRE to POST (within-SWR PRE reactivation mean: 1.7, median: 0.25, POST mean: 6.38, median: 1.85, Signed-Rank $p \sim 0$, $n = 3{,}371$ ensembles) and **(F)** this effect was consistent across mice. **(G)** Peri-SWR transient-based ICA ensemble activity was normalized as in fig. S10E, confirming **(H)** the within-SWR increases in RUN-template specific ensemble reactivation, **(I)** an effect which was consistent across animals. **(J-N)** Finally, an analysis paralleling that shown in Fig 4B, C, D, G and fig. S13A, respectively, was performed using the transient-based calcium activity estimates, further confirming that PRE to POST changes in SWR-specific recruitment predict future place cell stability and that this effect is specific to the path zone-coding cells.

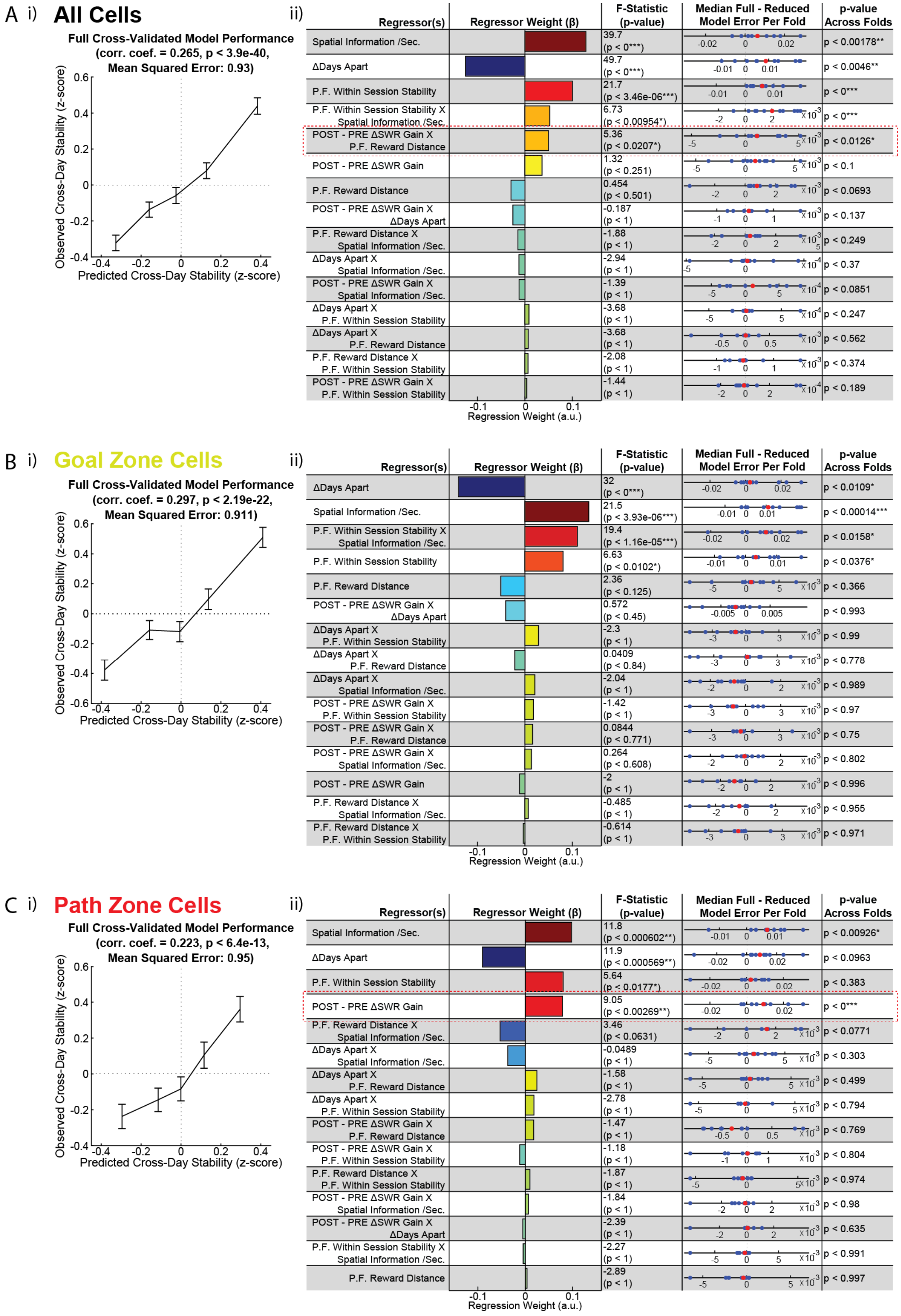


**Figure S15: Generalized linear model analysis of factors predictive of future stability.** In order to further test which factors predict future place field stability a 10-fold cross-validated ridge regression-based generalized linear model (GLM) was constructed. The model predicted future per-cell stability for novel place cells based on five unique factors normalized by session: 1) the cell's spatial information per second on the novel place field day, 2) the mean days apart between the novel place field day and the future days on the same RUN belt with which stability was compared, 3) the cell's normalized within session stability (z-scored –Δcm between the first and second half of laps on the novel place field day), 4) the cell's PF peak distance to reward on the novel place field day, and 5) the PRE to POST change in within-SWR gain observed on the novel place field day. In addition, the interactions between the 5 factors were also used as regressors, for a total of 15 regressors. **(A,** panel i**)** The GLM was found to be significantly predictive of future place field stability (plot shows the mean ±SEM of the observed normalized cross-day stability broken up into 6 equal parts by predicted cross-day stability (x-axis), Pearson's correlation between observed and predicted stability scores (r): 0.265, Fisher's z-test $p < 3.9 \times 10^{-40}$, n = 2,424 novel place cells). A subtractive analysis was performed by comparing the full-model GLM to reduced GLM's in which each of the 15 regressors were held out one at a time (all models 10 fold cross-validated). The table in panel ii shows the results of this subtractive analysis for the 15 regressors (left column), arranged from highest to lowest absolute regressor weight in the full model (second column). Reduced models were tested against the full model via two complementary methods: 1) a p-value was computed from the F-statistic of the difference in the mean squared error between the full and reduced models, 2) the difference in median absolute errors between the full vs reduced models was computed within each fold (fourth column, n = 10 folds, red markers show the cross-fold medians), and the significance of the difference in median errors was tested across folds (last column, p – value from one-sided boot-strap test of the mean, n = 10 folds). As expected, the quality of the place fields on the novel place field day as measured by the normalized spatial information content (first row) or within-session stability (third row), as well as the length of the temporal interval across which stability was assessed (Δdays apart, second row) were found to strongly predict future place field stability. That is, place cell's which were found to be more strongly place coding on their novel place field day tended to retain their place coding in the future and overall stability decayed over time (Fig. 1, I). However, after accounting for these expected effects via the subtractive GLM analysis and consistent with our other results (Fig 4, A-G) the interaction between novel place field day PRE to POST change in SWR-gain and P.F. reward distance was found to be significantly predictive of future place field stability (fifth row, dashed red box). **(B)** The GLM analyses were carried out after restricting the data to goal-zone cells (plotted as in panel A, n = 1,026 novel place cells). As expected, for this sub-group neither PRE to POST changes in SWR-gain or their interaction with P.F. reward distance were found to be significantly predictive of future place field stability. **(C)** By contrast, in path-zone cells (plotted as in panel A, n = 1,018 novel place cells) the PRE to POST change in SWR-gain regressor was found to significantly predict future place field stability (fourth row, dashed red box). These results confirm that after controlling for other factors expected to influence cross-day P.F. stability, consistent with POST epoch-dependent memory consolidation PRE to POST changes in SWR-gain, and their interaction with P.F. reward distance, are significantly predictive of future place field stability.

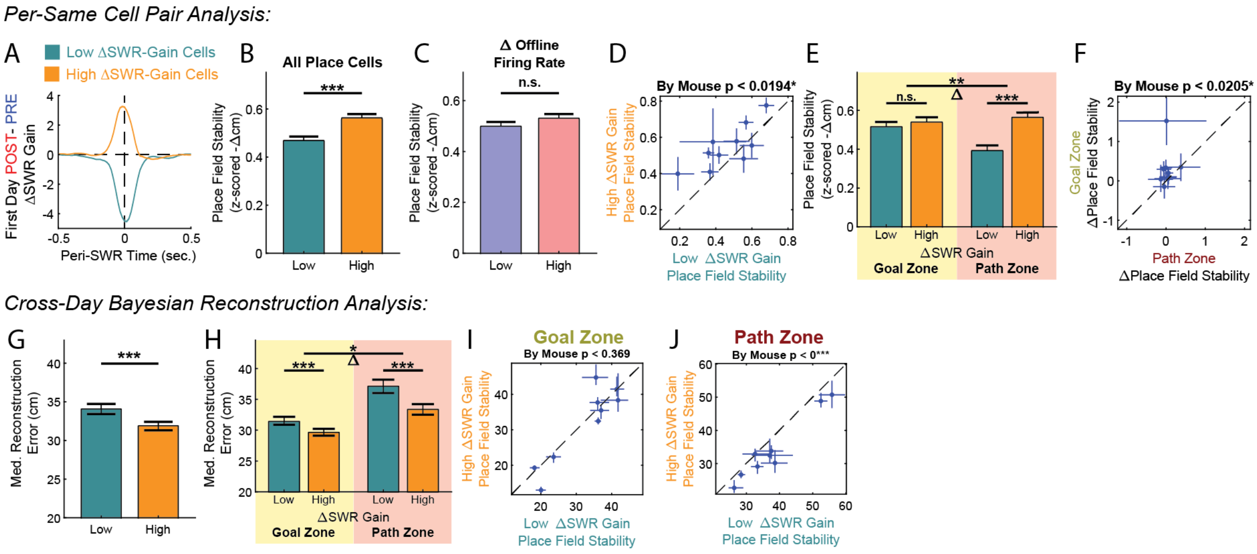


**Figure 16: 'Per same cell pair' and cross-day Bayesian reconstruction analysis.** In a complementary analysis to the 'Per cell' analysis carried out in Fig 4A-G a more inclusive 'Per same-cell pair analysis' was conducted in which each pair of times a given cell was a place cell on the same belt was considered separately (see Online Methods). **(A)** Same cell pairs were divided by their PRE to POST change in within-SWR gain on the earlier day of the pair (graphs show mean ±SEM peri-SWR firing rate gain for each of the two groups, n = 3,967 same-cell pairs per group). **(B-F)** PRE to POST changes in SWR-specific recruitment were found to predict future representational stability across days in the 'per same cell pair' analysis, and this effect was found to be larger in the Path as compared to Goal zones (graphs plotted as in fig. S13). The per-same cell pair groups shown in A-F were used for cross-day Bayesian reconstruction with earlier days serving as templates for location decoding of subsequent days on the same belt. **(G)** Graph shows the median decoding error across laps using either the Low or High ΔSWR-Gain groups (median ±boot-strapped 95% confidence interval of the median, Ranked-Sum test, n = 3,969 cross-day decoded laps in each group). Note the lower cross-day decoding error for the High ΔSWR-Gain group. **(H)** This analysis was then performed on the Goal and Path zones independently, revealing that while High ΔSWR-Gain cells were associated with improved decoding in both areas, the Low to High ΔSWR-Gain decoding error improvement was larger in the Path zone (Goal vs Path permutation test for interaction). **(I)** A per mouse analysis revealed that the Low vs. High ΔSWR-Gain group effect was not consistent across mice in the Goal zone (graph shows by mouse medians ±boot-strapped 95% confidence interval of the median, one of the mice did not have enough cross-day pair place cells in each of the groups (minimum 5 place cells per session pair, per cell group) to be included in the analysis, consequently, n = 9 mice, p-value from the one-sided boot strap test of the mean difference across animals). **(H)** However, mice consistently showed improved future-day location decoding using the High as compared to Low ΔSWR-Gain cells (n = 9 mice).

**Extended Data References:**
